# *De novo* assembly of a Tibetan genome and identification of novel structural variants associated with high altitude adaptation

**DOI:** 10.1101/753186

**Authors:** Ouzhuluobu, Yaoxi He, Haiyi Lou, Chaoying Cui, Lian Deng, Yang Gao, Wangshan Zheng, Yongbo Guo, Xiaoji Wang, Zhilin Ning, Jun Li, Bin Li, Caijuan Bai, Baimakangzhuo, Gonggalanzi, Dejiquzong, Bianba, Duojizhuoma, Shiming Liu, Tianyi Wu, Shuhua Xu, Xuebin Qi, Bing Su

**Author notes:** Corresponding author (B.S.); (X.Q.); (S.X.). These authors contributed equally to this work (Ouzhuluobu, Yaoxi He, Haiyi Lou, Chaoying Cui, Lian Deng).

## Abstract

Structural variants (SVs) may play important roles in human adaption to extreme environments such as high altitude but have been under-investigated. Here, combining long-read sequencing with multiple scaffolding techniques, we assembled a high-quality Tibetan genome (ZF1), with a contig N50 length of 24.57 mega-base pairs (Mb) and a scaffold N50 length of 58.80 Mb. The ZF1 assembly filled 80 remaining N-gaps (0.25 Mb in total length) in the reference human genome (GRCh38). Markedly, we detected 17,900 SVs, among which the ZF1-specific SVs are enriched in GTPase activity that is required for activation of the hypoxic pathway. Further population analysis uncovered a 163-bp intronic deletion in the *MKL1* gene showing large divergence between highland Tibetans and lowland Han Chinese. This deletion is significantly associated with lower systolic pulmonary arterial pressure, one of the key adaptive physiological traits in Tibetans. Moreover, with the use of the high quality *de novo* assembly, we observed a much higher rate of genome-wide archaic hominid (Altai Neanderthal and Denisovan) shared non-reference sequences in ZF1 (1.32%-1.53%) compared to other East Asian genomes (0.70%-0.98%), reflecting a unique genomic composition of Tibetans. One such archaic-hominid shared sequence, a 662-bp intronic insertion in the *SCUBE2* gene, is enriched and associated with better lung function (the FEV1/FVC ratio) in Tibetans. Collectively, we generated the first high-resolution Tibetan reference genome, and the identified SVs may serve as valuable resources for future evolutionary and medical studies.

## Introduction

Next generation sequencing (NGS) is a powerful tool to study human genomic variations through simple alignment of short reads to a reference genome. However, short reads have unavoidable limitations for genome assembly, especially for detection of structural variants (SVs) that have been shown to play an important role in normal and abnormal human biology^1, 2^. By contrast, with an advantage of long reads (>10 kilo-base pair, kb), the single-molecular real-time (SMRT) sequencing (also called the third-generation sequencing, TGS) has been proven effective in resolving complex genomic regions, such as sequences with SVs^3, 4^. Meanwhile, the application of next generation mapping technologies provides complementary approaches to *de novo* genome assembly, including BioNano, 10X Genomics and Hi-C etc. Recently, with the aid of SMRT sequencing and next generation mapping methods, two long-read Asian genome assembles (AK1 and HX1) were released^5, 6^.

Tibetans represent a unique highland population permanently living at the Tibetan Plateau (average elevation >4,000 meters), one of the most extreme environments on earth. Their permanent settlement in the Qinghai-Tibetan plateau was dated as early as 30,000 years ago based on genetic data (Shi, et al. 2008; Qi, et al. 2013; Lu, et al. 2016). Previous genetic studies have identified two key genes (*EPAS1* and *EGLN1*) carrying adaptive alleles that help maintain relatively lower hemoglobin concentration in native Tibetans so that over-production of red cells (polycythemia) at high altitude could be avoided ^7–17^. Also, a Tibetan-enriched 3.4kb deletion (TED) near *EPAS1* was reported^18^. Additionally, it was proposed that the Tibetan-enriched *EPAS1* variants were inherited from Denisovan-like hominid^19^. These evidences suggest that the high-altitude adaptation of Tibetans is probably multi-facet, involving different types of genomic variations.

Besides hemoglobin concentration, there are other key adaptive physiological traits in Tibetans, such as elevated resting ventilation and low hypoxic pulmonary vasoconstrictor response^20^, which cannot be explained by the known single nucleotide variations (SNVs) identified using NGS data. Putatively, SVs located in the regulatory regions of the genome may contribute to these unresolved adaptive traits. Also, the sequences present in Tibetans but absent in the human reference genome are putative introgressions from archaic humans, which have not been systematically evaluated. Hence, these unsolved questions call for a high-quality Tibetan reference genome.

We combined SMRT long-read sequencing with multiple scaffolding techniques, as well as short-read deep-sequencing, and we *de novo* assembled a high-quality Tibetan genome (ZF1). The assembled Tibetan genome reached a contig N50 size of 24.57Mb and a scaffold N50 size of 58.80Mb. We used a read-mapping approach to detect SVs in the assembled ZF1 genome. By comparing with two previous long-read Asian genome assemblies (AK1 and HX1), we identified a large number of novel SVs, some of which are enriched in Tibetans and showed association with pulmonary arterial pressure and lung functions. Furthermore, using the high quality ZF1 assembly, we found a much higher rate of genome-wide archaic hominid (Altai-Neanderthal and Denisovan) shared non-reference sequences in ZF1 than in other East Asian genomes.

## Results

### De novo assembly of the Tibetan genome and gap filling on the reference genome

We performed SMRT long-read sequencing using PacBio RSII at 70× coverage and obtained a total of 24.9M subreads with median and mean read-length of 9.5kb and 10.3kb, respectively (Supplementary Figure 1). The long reads were error-corrected and assembled into contigs by Falcon, and then the assembled contigs were polished by Quiver^21^. In total, we generated 3,148 contigs with a N50 length of 23.62Mb (Supplementary Figure 2 and Supplementary Tables 1-3, Methods). To order and link these contigs into larger scaffolds, we utilized the data of BioNano and 10X Genomics, and constructed two versions of scaffolding (Supplementary Table 2, 3). The first version used the 10X Genomics linked-reads (100×) to link the contigs into larger scaffolds, and then combined the physical maps with unique motifs from BioNano. This scaffolding strategy resulted in 2,403 scaffolds with a N50 length of 45.42Mb (Supplementary Figure 2a and Supplementary Table 2, Methods). The second version used the BioNano data first and then the 10X Genomics reads, which resulted in 2,321 scaffolds with a N50 length of 47.17 Mb (Supplementary Figure 2b and Supplementary Table 3). Considering the longer scaffold N50 length, we chose the second version for further improvement using Hi-C data.

We generated 100× Hi-C data of ZF1 (Supplementary Table 1). Using SALSA^22^, we grouped the scaffolds using Hi-C data, leading to a final version of the assembled ZF1 genome of 2.89 Gb with a contig N50 length of 24.57Mb and a scaffold N50 length of 58.80Mb. In addition, we performed short-read sequencing using Illumina HiSeq X10 platform and generated 100× coverage of the ZF1 genome to improve base-level accuracy (Figure 1 and Supplementary Table 2, 3, Methods). We also generated a phased version of the ZF1 genome assembly (see Methods for technical details).

**Figure 1.**
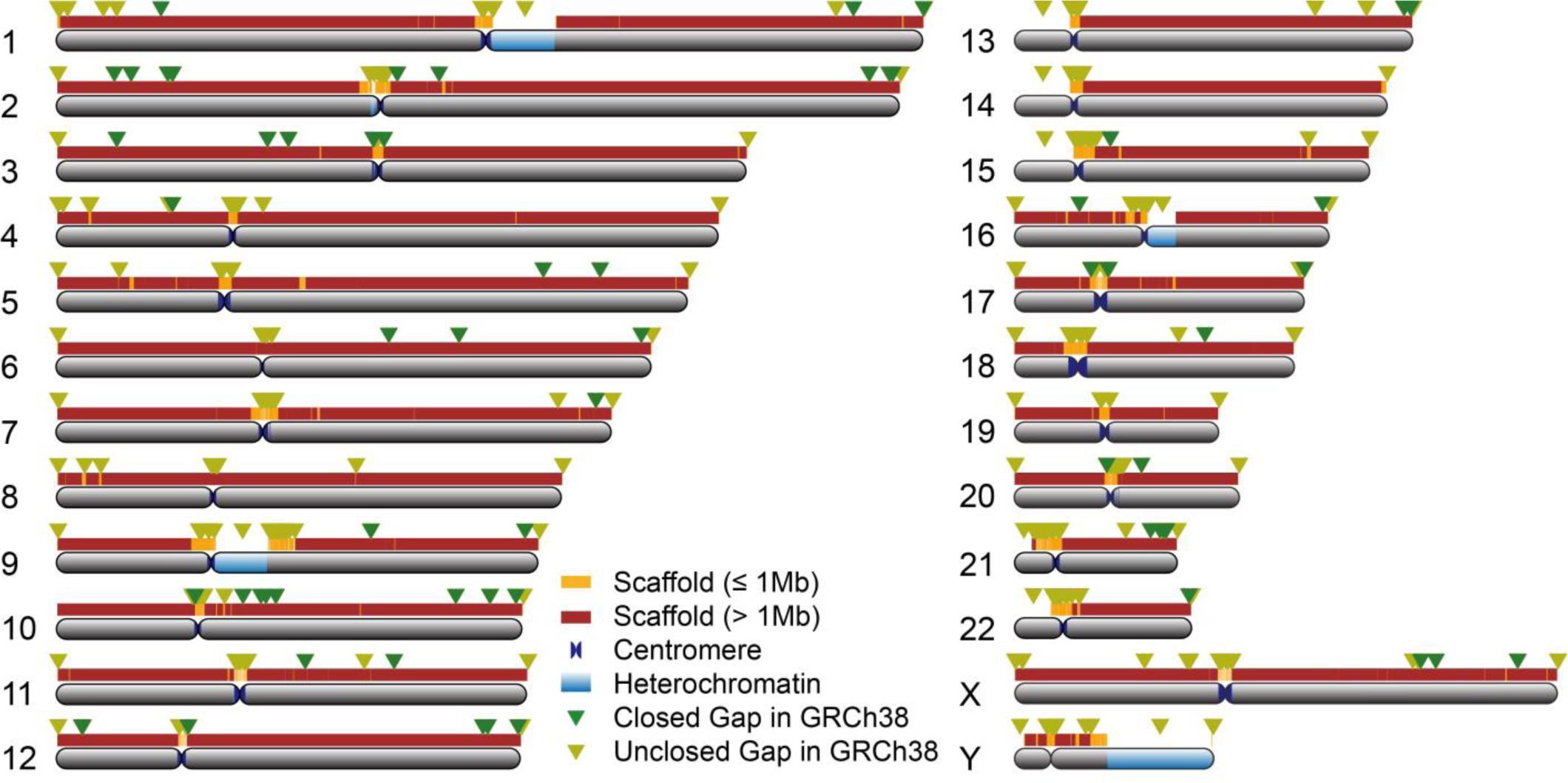
*De novo* assembly of the ZF1 genome compared to GRCh38. Scaffold coverage and gap closure over GRCh38 per chromosome are shown in the plot. The colored bars above each chromosome represent the ZF1 scaffolds, with the dark red segments for the long scaffolds (> 1Mb) and the orange segments for the short ones (≤ 1Mb). Closed euchromatic gaps are labeled by the green triangles on each chromosome, and the unfilled gaps in ZF1 by the red triangles.

Next, we used the *de novo* assembled ZF1 genome to conduct gap closure for the human reference genome GRCh38. A total of 80 of the 940 N-gaps in the GRCh38 human reference genome were completely filled by the ZF1 assembly, and the total length was 0.25Mb (Figure 1, Table 1 and Supplementary Table 4, Methods).

**Table 1.** Comparison of the ZF1 *de novo* assembly with the published human *de novo* assemblies.

| Assembly | Assembly approach | Sequencing platform | Contig N50 (Mb) | Scaffold N50 (Mb) | Genome size (Gb) | Gaps number | Gap length (Mb) |
| --- | --- | --- | --- | --- | --- | --- | --- |
| ZF1 | WGS, Hi-C | PacBio, BioNano, 10X Genomics, Hi-C, Illumina-PE | 24.57 | 58.8 | 2.90 | 740 | 7.82 |
| HX1 | WGS | PacBio, BioNano, Illumina-PE | 8.33 | 21.98 | 2.93 | 10901 | 39.34 |
| AK1 | WGS, BAC | PacBio, BioNano, Illumina-PE | 17.92 | 44.85 | 2.90 | 264 | 37.34 |
| GRCh38 | BAC, Fosmid | Sanger, FISH, OM, fingerprint contigs | 56.41 | 67.79 | 3.20 | 940 | 159.97 |
| NA12878 (ASM101398) | WGS | PacBio, BioNano | 1.56 | 26.83 | 3.20 | 2332 | 146.35 |

WGS: Whole genome sequencing; BAC: Bacterial artificial chromosome; PE: Pair end; FISH: fluorescence in situ hybridization; OM, optical mapping.

To evaluate the completeness and accuracy of the ZF1 genome assembly, we compared the ZF1 assembly with two previous long-read Asian genome assemblies (AK1^5^ and HX1^6^) and a high-quality European genome (ASM101398, sample ID: NA12878^23^. We found that the total bases (non-N bases in assembly) of the three Asian *de novo* assemblies (ZF1, AK1 and HX1) were quite similar (Table 1 and Supplementary Figure 3). Notably, the remaining gap length of ZF1 (7.82Mb) is much shorter than those in AK1 (37.34Mb) and HX1 (39.34Mb) (Supplementary Table 5). In addition, using MUMmer^24^, we assessed the consensus quality of the ZF1 assembly by aligning the ZF1 chromosomes with those of GRCh38, and we obtained 99.90% consensus accuracy for the ZF1 assembly, which is better than those for HX1 (99.73%), YH2.0 (99.81%), NA12878 (99.73%) and HuRef (99.84%)^6^ (Supplementary Figure 4). Additionally, we evaluated the base-error rate of the ZF1 assembly using our 100× Illumina short-read data with the previous approach^25, 26^. The inconsistency rate is 0.0006% (Supplementary Table 6), well below one error per 10,000 bases, the quality standard used for human genome^27^.

Furthermore, we annotated the ZF1 assembly using CESAR2.0^28^ and made functional annotation for ZF1 genes with four databases (KEGG, Swiss-Prot, InterPro and NR) (Supplementary Figure 5). We obtained similar number of annotated genes compared with GRCh38 (ZF1: 19,805 vs. GRCh38: 19,267), and 99.8% of the ZF1 genes were annotated by multiple databases. Notably, the ZF1 assembly embraced a longer average CDS length than GRCh38 (Supplementary Figure 6 and Supplementary Table 7, 8), and this improvement may stem from long-read assembly of more exons of the ZF1 assembly. Taken together, our ZF1 assembly provided a reliable reference genome for downstream analysis.

### Profiling SVs of the ZF1 genome

We employed a read-mapping-based approach to call SVs from the ZF1 PacBio long reads (see Methods for details) ^4^. For the compatibility of conducting downstream SV comparison analysis with the public data, we used the human genome build GRCh37 instead of GRCh38 as the reference. Within the size range of 50bp to 2Mb, we obtained 17,900 SVs, including 7,461 deletions, 1,853 duplications, 8,196 insertions, 204 inversions and 186 complex SVs, among which 75% of the SVs were supported by results from other platforms (*i.e.* Illumina X10, BioNano and 10X Genomics; Figure 2a, Supplementary Table 9 and 10). The distribution of the large SVs (>1kb) on the 24 chromosomes (including X and Y chromosomes) of ZF1 is shown in Supplementary Figure 7. The median lengths for deletion, insertion, duplication and inversion are 166bp, 144bp, 543bp and 1,399bp, respectively. These SVs cover 57.8 Mb in total, accounting for ∼2% of the entire genome (Figure 2b). Almost 70% of the SVs contain repetitive elements such as SINEs, LINEs, simple repeats and satellites etc. (Supplementary Table 11).

**Figure 2.**
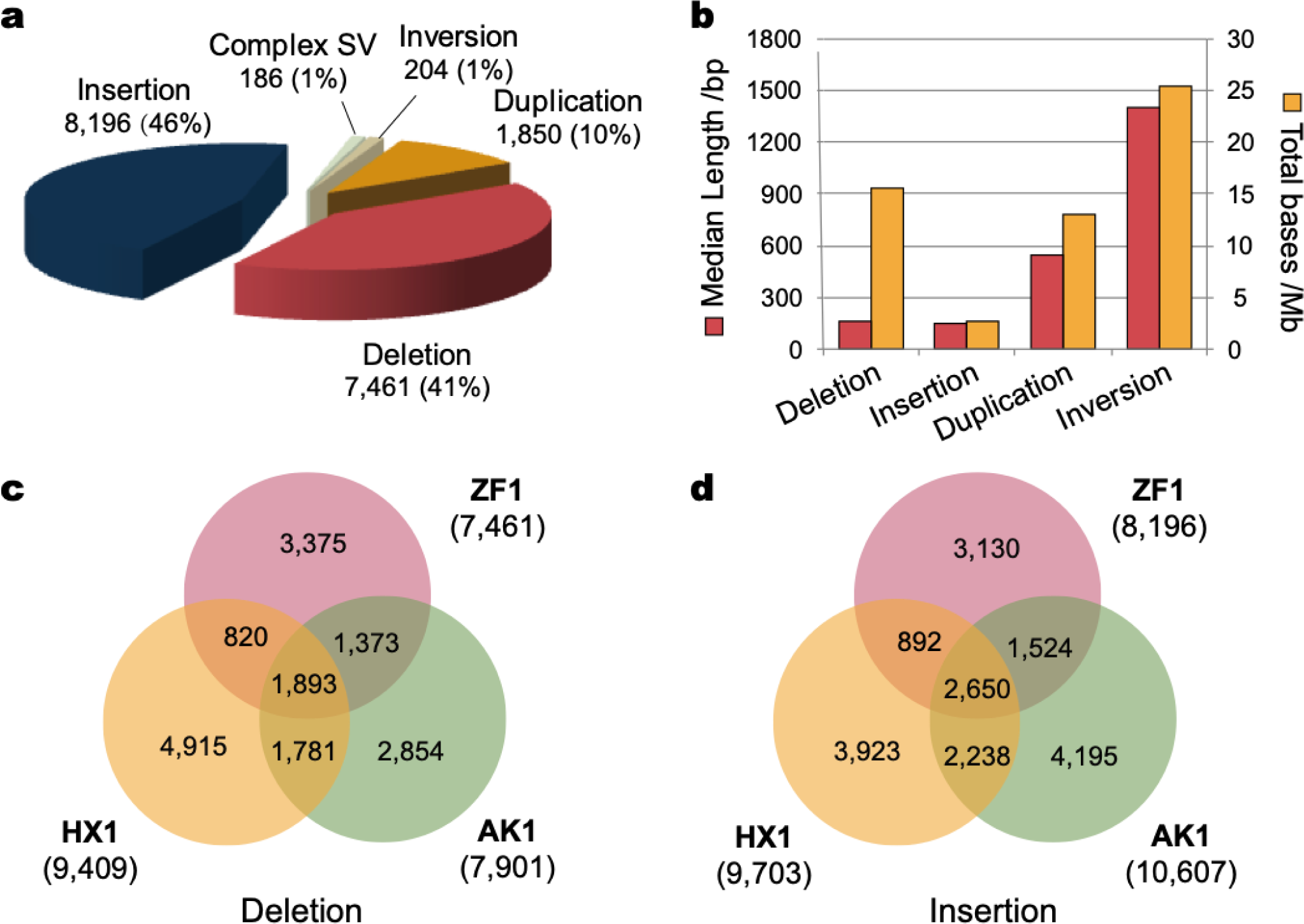
A summary of structural variants detected in ZF1. **a.** Pie plot shows the number and the proportion of insertions, deletions, duplications, inversions and complex SVs detected in ZF1. **b.** Median length and total base statistics of different SVs. **c, d.** Overlap of SVs among ZF1, HX1 and AK1 for deletions (**c**) and insertions (**d**), respectively. An overlapped SV was defined as one with overlapping length reaching at least 50% of reciprocal similarity.

Compared with AK1 and HX1, ZF1 has 6,505 specific SVs (36.3%, 6,505/17,900) (the SVs not found in either AK1 or HX1), including 3,375 deletions and 3,130 insertions, accounting for 45.24% (3,375/7,461) and 38.19% (3,130/8,196) of the total deletions and insertions, respectively (Figure 2c, 2d). We annotated these ZF1-specific SVs to their nearby genes (within 5kb downstream or upstream of the SVs) using VEP (Variant Effect Predictor). Totally, we found 1,832 genes near the ZF1-specific SVs, and we defined them as ZF1-specific-SVs-associated genes (ZSAGs). Functional enrichment analysis showed that these ZSAGs were enriched in four functional clusters, *i.e.* positive regulation of GTPase activity (false discovery rate, FDR=1.78E-05), intracellular signal transduction (FDR=8.03E-03), transmembrane receptor protein tyrosine kinase signaling pathway (FDR=1.07E-02) and peptidyl-tyrosine phosphorylation (FDR=3.47E-02) (Supplementary Figure 8 and Supplementary Table 12).

We next explored how many ZSAGs were related to hypoxic regulation. Among the 571 priori candidate genes of hypoxia adaptation in the Tibetans (473 known hypoxia-related genes ^16^ and 168 reported genes showing signals of Darwinian positive selection in Tibetans^7–12, 14, 29^), we found 69 of them overlapped with the 1,832 ZSAGs (Supplementary Table 13-14; odds ratio = 2.42, *p* = 3.4 × 10^-12^, Chi-squared test). Interestingly, these newly identified hypoxia- and selection-related SVs are all located in either intronic or intergenic regions, and if functional, they are more likely to affect gene expression regulation.

### Novel Tibetan-specific SVs are associated with high-altitude adaptation

The high-resolution ZF1 genome assembly provides high confidence SVs with precise breakpoint positions, which could serve as a reference panel to assess the variant frequency difference between Tibetans and lowland populations. To this end, we analyzed the genetic divergence of SVs between Tibetans and Han Chinese using the published population NGS data (38 Tibetans and 39 Han Chinese, see Methods for details). We listed the 124 SVs (the top 5% of the 1,887 CNVs and 593 insertions that passed the NGS-genotyping filtering; see Methods) with the highest between-population divergence (measured by VST) in Supplementary Table 15 and 16. The most diverged SV was the previously reported TED near *EPAS1* (VST = 0.725)^18^(Supplementary Figure 9 and Supplementary Table 15). The remaining 123 SVs contain 93 copy number variations (Supplementary Figure 9 and Supplementary Table 15) and 30 insertions (Supplementary Figures 10 and Supplementary Table 16).

Notably, we found a 163bp-deletion (chr22:40,935,468-40,935,631, hg19) with an allelic divergence of 0.227 between Tibetans and Han Chinese (the allele frequencies are 0.544 and 0.317 in Tibetans and Han Chinese, respectively) (Supplementary Table S17). The VST of this variant (0.106) is among the top 5% of the genome-wide SVs (5% cut-off = 0.0956), a suggestive signal of selection and less likely caused by genetic drift or other demographic events according to the simulation analysis (see Methods, Supplementary Figure 19).

This deletion is located in the intronic region of *MKL1*, which encodes Megakaryoblastic Leukemia 1 and was previously reported to regulate hypoxia induced pulmonary hypertension in rodents^30, 31^. With the use of the ENCODE data, we found multiple histone modification signals overlapped with the 163bp-deletion, suggesting that this SV is located in a region with enhancer activity. We also detected a GeneHancer regulatory element (MKL1/GH22J040443) and two methylation hotspots in this region (Figure 3a and Supplementary Figure 11).

**Figure 3.**
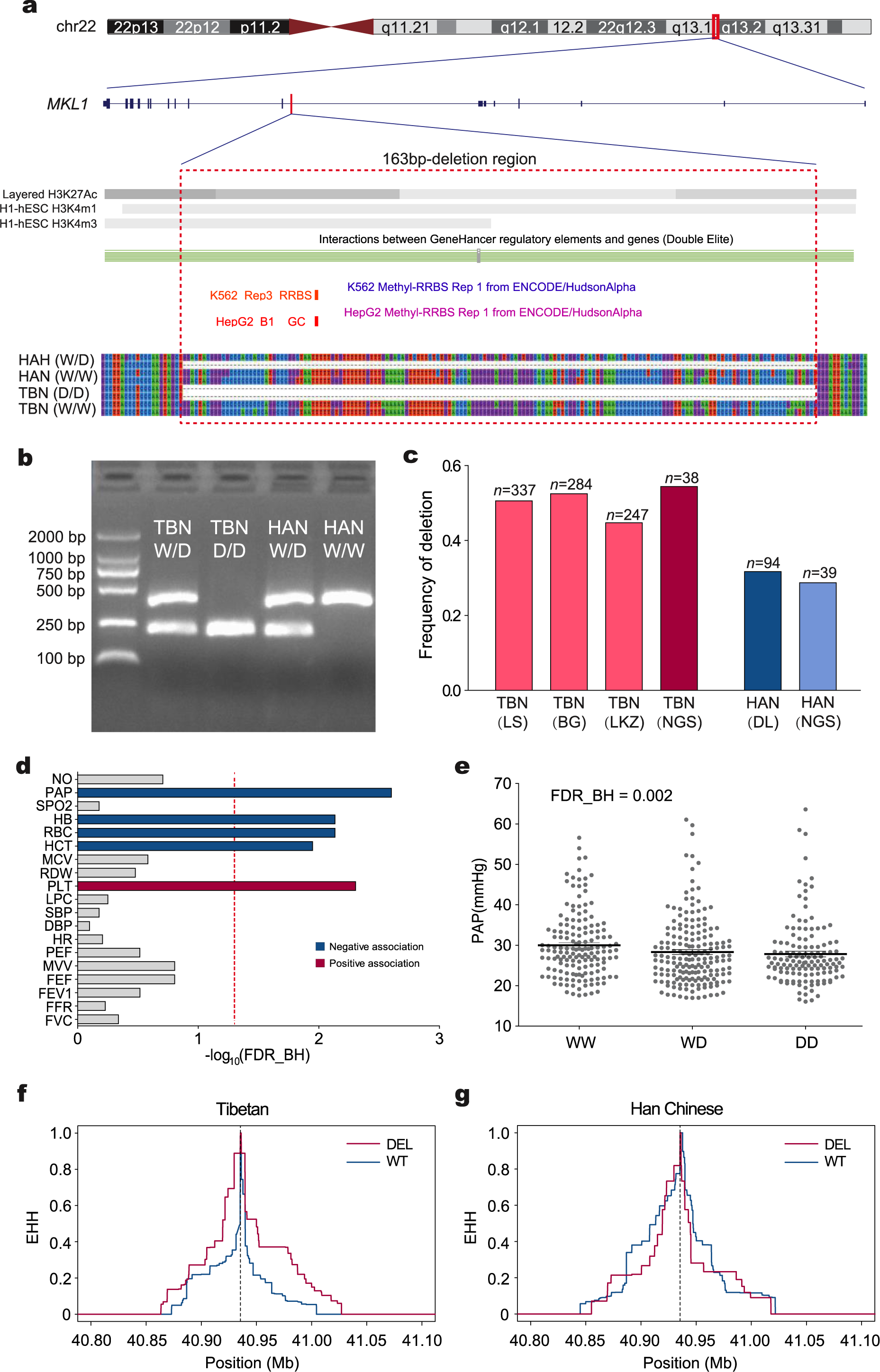
The Tibetan-enriched *MKL1* 163bp-deletion and its association with physiological traits. **a.** The schematic map indicating the genomic location (upper panel), epigenetic signals (histone modification and DNA methylation, middle panel) and sequence alignment (bottom panel) of the *MKL1* deletion and its flanking sequences in Tibetans and Han Chinese; **b.** Genotyping electromorphic of the *MKL1* deletion. The two alleles are indicated as “W” (wildtype) and “D” (deletion); **c.** Allele frequencies of the *MKL1* deletion in Tibetans (TBN) and Han Chinese (HAN); TBN(LS): Tibetans at Lhasa, TBN(BG): Tibetans at Bange, TBN(LKZ): Tibetans at Langkazi, HAN(DL): Han Chinese at Dalian; TBN(NGS) and HAN(NGS): Tibetans and Han Chinese from the NGS data (Methods); **d.** Genetic association between the *MKL1* deletion and multiple physiological traits in Tibetans (n=868). The dot line in red refers to the cutoff of statistical significance with false discovery rate of less than 5% by Benjamini & Hochberg (FDR_BH)^69^, and the trait abbreviations are described in Methods. **e.** Comparison of pulmonary arterial pressure levels among three different genotypes at the *MKL1* deletion; W-wide type (non-deletion). P value was calculated assuming an additive model with multiple testing correction using Bejamini & Hochberg FDR control (Methods). **f-g.** Estimation of EHH decay of haplotypes in Tibetans (f) and Han Chinese (g) surrounding the *MKL1* 163bp-deletion. The physical position of the 163bp-deletion was indicated by the vertical dashed line in (f) and (g).

We measured 19 physiological traits (varied blood, heart and lung indexes) and collected blood samples from 1,039 indigenous adult Tibetans. Using PCR (polymerase chain reaction) and Sanger sequencing, we genotyped the *MKL1* deletion in 868 Tibetans from three geographic populations, including 337 unrelated individuals from Lhasa (elevation: 3,658m), 284 unrelated individuals from Bange (elevation: 4,700m) and 247 unrelated individuals from Langkazi (elevation: 5,108m). We also genotyped 94 unrelated Han Chinese from northern China (elevation: 60m). The frequencies of the *MKL1* deletion are similar with those estimated based on NGS data (0.500 in Tibetans and 0.287 in Han Chinese; Figure 3c and Supplementary Table 17, Methods). We next performed association analysis in Tibetans. Since no genetic heterogeneity was detected among the three Tibetan populations, the raw data was merged (n=868). We found that the *MKL1* deletion was negatively associated with PAP (systolic pulmonary arterial pressure) (FDR=0.002, Figure 3e), and the *MKL1* deletion carriers tend to have a lower PAP, consistent with the well-known low hypoxic pulmonary vasoconstrictor response in Tibetans^20^. Interestingly, the *MKL1* deletion also showed association with several blood indices, including negative associations with HB (hemoglobin concentration) (FDR=0.03), HCT (hematocrit) (FDR=0.02) and RBC (red blood count) (FDR=0.01), and a positive association with PLT (platelets) (FDR=0.004) (Figure 3d and Supplementary Table S18), implying that the *MKL1* deletion might be involved in multiple regulatory effects on pulmonary and blood indexes. When using the most rigorous adjustment (19 traits × 4 SVs) for multiple test correction, PAP, RBC and PLT still remain significant: FDR(PAP)=0.008, FDR(RBC)=0.04 and FDR(PLT)=0.008.

In addition to the *MKL1* deletion, we also selected other SV overlapping genes with previous evidence of positive selection or related with hypoxia regulation (Supplementary Table 19). Although these SVs were not among the top 5% Tibetan-Han diverged SVs, some of them were significantly associated with multiple physiological traits (Supplementary Table 18). For example, a 53bp-insertion (allele frequency: 0.424 in TBN and 0.339 in HAN, Supplementary Table 19) in *COL6A2* was significantly associated with systolic blood pressure (SBP) and diastolic blood pressure (DBP) (*P*=0.0023, FDR=0.03; *P*=0.0054, FDR=0.035, respectively; Supplementary Figure 12, 13 and Supplementary Table 18). *COL6A2* encodes eukaryotic translation initiation factor 4E with selection signals in Ethiopian high-altitude population^32^. Another example is a 63bp-insertion in *EIF4E2*. The protein encoded by *EIF4F2* can form a complex with HIF-2 (encoded by *EPAS1*) and RBM4 under hypoxia as an oxygen-regulated switch^33^. This insertion (allele frequency: 0.457 in TBN and 0.382 in HAN) was significantly associated with maximum ventilatory volume (MVV) (*P*=0.04, FDR=0.293, Supplementary Figure 12 and Supplementary Table 18).

### Identification of ZF1-specific novel sequences shared with archaic humans

Previous studies have found evidence of Denisovan-like archaic introgression in the Tibetan genome such as the 32.7 kb fragment in *EPAS1*^19^ and a ∼300 kb region in the chromosome 2 derived from unresolved archaic ancestry^34^. However, previous results were mainly inferred from SNVs with respect to the human reference genome. The sequences present in both archaic and Tibetans but absent in the human reference genome have not been systematically assessed. To take advantage of the *de novo* ZF1 assembly, we performed a genome-wide search of archaic sharing non-reference sequences (NRSs) and compared the results with the two *de novo* assembled Asian genomes (AK1 and HX1) (Methods). We found a total length of 39.6Mb and 45.9Mb sequences shared with those of Altai Neanderthal and Denisovan, corresponding to 1.32% and 1.53% of the entire ZF1 genome respectively. These archaic proportions are much higher than that in AK1 (0.82% and 0.70%) or HX1 (0.98% and 0.85%) (Figure 4a). We further checked those novel archaic-shared sequences that could be unambiguously determined as an insertion model (Methods), and identified 239 Neanderthal- and 164 Denisovan-shared ZF1-specific events (sequences only present in ZF1 but absent in AK1 or HX1), in contrast to the 133 Neanderthal- and 115 Denisovan-shared AK1-specific, and 151 Neanderthal- and 126 Denisovan-shared HX1-specific events, indicating the Tibetan genome contains more archaic-shared sequences than the other two East Asian genomes. After further filtering using the published European (NA12878) and African (NA19240) genomes, we obtained 167 Neanderthal- and 117 Denisovan-shared ZF1-specific events that were absent in the representative modern human assemblies (Methods, Figure 4b, Supplementary Table 20), among which 51 Neanderthal- and 28 Denisovan-shared ZF1-specific events are present in great apes (chimpanzee, gorilla and orangutan).

**Figure 4.**
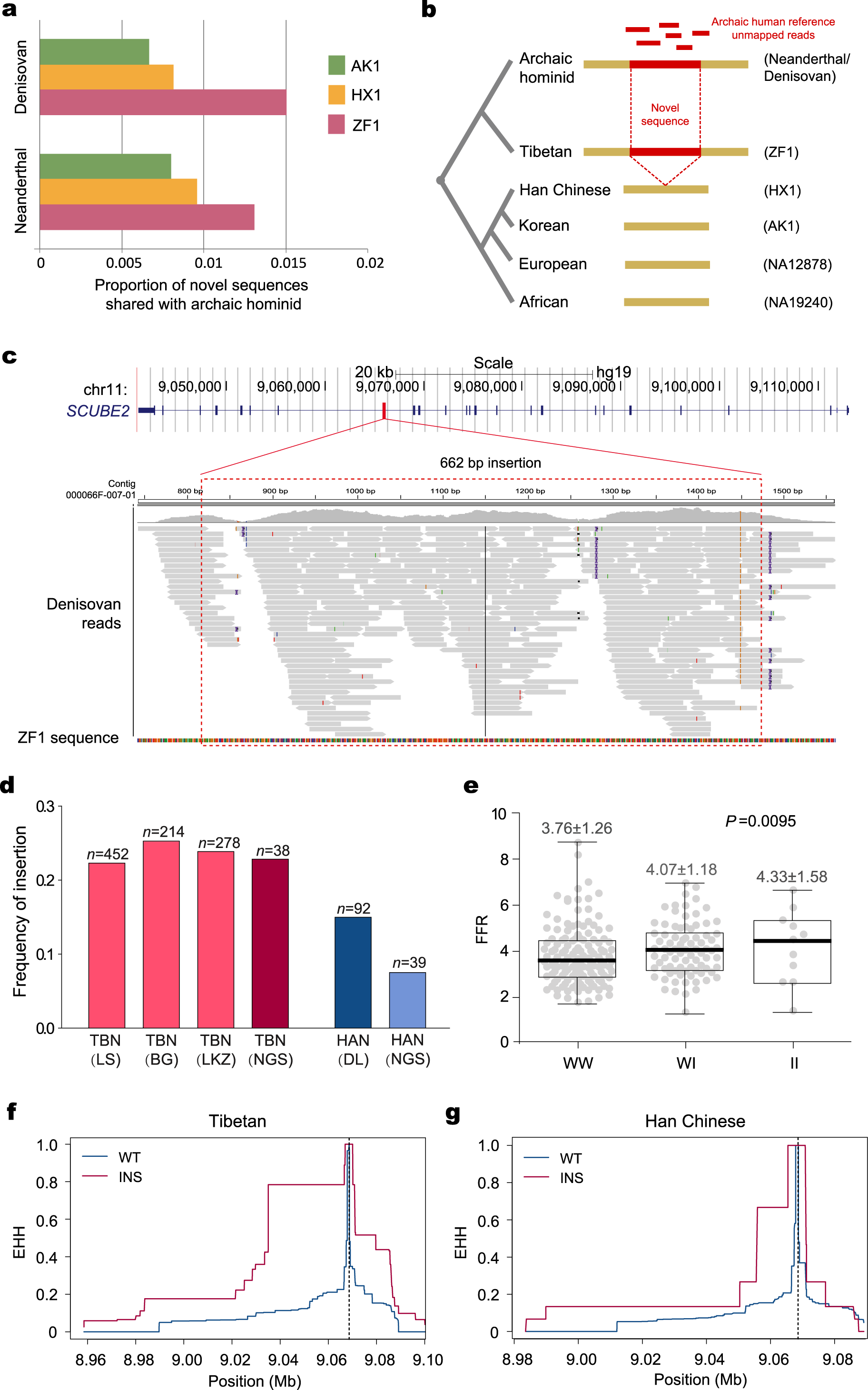
ZF1-speicific novel sequences shared with archaic hominins. **a.** Genomic proportion of the non-reference sequences shared with archaic hominids in the three *de novo* assembled Asian genomes. **b.** Schematic diagram of the ZF1-sepcific non-reference sequences shared with archaic hominids. Novel non-reference sequences (red) shared between Tibetan (ZF1) and archaic hominids but absent in the other representative modern human genomes (one European, one African and two Asian genomes). **c.** A novel sequence shared between ZF1 and archaic hominid in the intron of *SCUBE2*. This sequence was found in both Denisovan and Altai Neanderthal. Shown here is the Denisovan reads mapping. The upper panel shows the position of this 662-bp novel sequence in the human reference genome GRCh37. The bottom panels indicate the Denisovan reference unmapped reads aligned to the ZF1 contig. **d.** Allele frequencies of the 662bp-insertion in Tibetans (TBN) and Han Chinese (HAN), TBN(LS): Tibetans at Lhasa; TBN (BG): Tibetans at Bange; TBN(LKZ): Tibetans at Langkazi; HAN(DL): Han Chinese at Dalian; TBN(NGS) and HAN(NGS): insertion frequency estimated by next-generation-sequencing data (Methods). **e.** Comparison of the FEV1/FVC ratios among three different genotypes at the *SCUBE2* insertion; I-insertion; W-wide type (non-insertion). **f-g.** Estimation of EHH in TBN (f) and HAN (g) around the *SCUBE2* insertion at Chr11:9068607. The physical position of the insertion was indicated by the vertical dashed line in (f) and (g).

Among the archaic-shared ZF1-specific NRSs, we found a 622-bp sequence in the intron of *SCUBE2* (Signal peptide-, CUB domain-, and EGF-like domains-containing protein 2) (Figure 4c), a non-repetitive insertion listed as the top3 diverged insertions between Tibetans and Han Chinese (mVST=0.079, Supplementary Figure 10 and Supplementary Table 16). The 622-bp *SCUBE2* sequence is also present in great apes, suggesting that it is an ancestral sequence. We genotyped the *SCUBE2* insertion using PCR and Sanger sequencing in the three Tibetan populations (452 Lhasa samples, 214 Bange samples and 278 Langkazi samples), as well as in the Han Chinese population (92 samples) (Supplementary Figure 14). The allele frequency of the *SCUBE2* insertion in Tibetans is on average near two-folds of that in Han Chinese (0.240 vs 0.130 for the combined samples, Figure 4d and Supplementary Table 21). The VST[CN] of this variant (0.096) is among the top 5% of the genome-wide SVs (5% cut-off = 0.0956), and is also significantly larger than the expected value under neutrality (see Methods, Supplementary Figure 19).

Previous study found that *SCUBE2* could regulate VEGF-induced angiogenesis^35^. To check the functional relevance of the *SCUBE2* 662-bp insertion, we performed genetic association analysis in Tibetans (n=944). We detected positive association with one lung index, the FEV1/FVC ratio (FFR) (FVC-forced vital capacity) (*P*=0.05, FDR=0.28) (Figure 4e and Supplementary Table 18), although the association became non-significant after multiple-testing correction. In addition, using a joint additive-model, we performed association analysis by combing the two SVs (the *MKL1* 163-bp deletion and the *SCUBE2* 662-bp insertion), and we observed a stronger signal for FFR compared with the single-SV analysis (*P*=0.00952, FDR=0.0602; Supplementary Figure 15). It was known that Tibetans perform better than Han Chinese in view of lung functions at high altitude (*i.e.* larger FVC and FEV1)^20, 36, 37^. Collectively, these results suggest that the two SVs may work together to improve lung function of Tibetans.

## Discussion

Through an integrated approach using PacBio long-read sequencing, BioNano optical mapping, 10X Genomics, Illumina HiSeq X10 and Hi-C technologies, we *de novo* assembled a high-quality Tibetan genome (ZF1). Compared with the previous *de novo* assemblies, the ZF1 assembly showed substantially improved quality with longer contig and scaffold N50 sizes. Based on this high-quality Tibetan genome, we detected 6,505 ZF1-specific SVs, and the associated genes are enriched for four functional clusters, especially for GTPase activity. Notably, GTPase activity is required for activation of hypoxia-inducible factor 1 (HIF-1α). In hypoxic cells, the small GTPase Rac1 is activated in response to hypoxia and is required for the induction of HIF-1α protein expression and transcriptional activity^38^. Consistently, the previously reported genes under selection in Tibetans (Supplementary Table 14) are enriched in the HIF-1 signaling pathway. Presumably, natural selection might have picked up some of these SVs contributing to high altitude adaptation in Tibetans. Further population and functional data are needed to test the contribution of the GTPase-activity-related SVs to the regulation of the hypoxic pathway.

The identified SVs with base-level breakpoints accuracy offer a comprehensive map that could facilitate estimating the variant frequencies in the corresponding populations so that SVs with large divergence between highlander Tibetans and lowlander Han Chinese can be found. Importantly, combining TGS and NGS data, we successfully identified an intronic 163bp-deletion in the intron of *MKL1* and a 662-bp insertion in *SCUBE2* that are highly differentiated between Tibetans and Han Chinese. We speculate that these two SVs are likely under positive selection in Tibetans: 1) the considerable genetic differentiation between highlander Tibetans and lowlander Han Chinese is less likely caused by genetic drift or other demographic events under neutrality according to our simulation data; 2) they show significant associations with multiple adaptive physiological traits in Tibetans, and might have larger functional influence than SNPs in terms of nucleotide length. We noted that the two SVs did not show significant iHS or XP-EHH estimates over the genome, but this should not be the reason to rule out the possibility of positive selection on these loci. It is well-acknowledged that different methods for detecting positive selection have their own underlying principles and weakness, and we should not expect positive results for a potential signal from all of them. In particular, most of the current methods were designed for SNP data, and could have limited power when applied to SV analysis. The substantial genetic differentiation and phenotypic association suggest weak selection on the deletion at *MKL1* and the insertion at *SCUBE2*. Functional validation of these novel SVs would largely rely on the experimental studies in the future.

*MKL1* is a transcriptional regulator known to influence cellular response to stress signals in the vasculature. It was shown that under chronic hypobaric hypoxia, the lung expression of *MKL1* was up-regulated in both rat and mouse, and *MKL1* knockdown could attenuate hypoxia induced pulmonary hypertension (HPH)^30, 31^. The *MKL1* protein directs histone H3 lysine 4 methyltransferase complexes to ameliorated HPH in mice^30^. Accordingly, the Tibetan-enriched *MKL1* 163bp-deletion is located in a putative enhancer sequence and embraced a GeneHancer regulatory element and two methylation hotspots (Figure 3a and Supplementary Figure 11), suggesting that it may affect epigenetic regulation of *MKL1* and eventually the downstream pathways, including vascular remodeling, vascular tone, and pulmonary inflammation. Consistently, we saw negative association of the *MKL1* deletion with pulmonary arterial pressure in Tibetans, explaining their low hypoxic pulmonary vasoconstrictor response at high altitude^20^. In line with this view, based on SNV analysis, *MKL1* was recently reported undergone positive selection in the Himalayan populations from Nepal, Bhutan, North India and Tibet^29^. The 163bp-deletion may disrupt an enhancer of *MKL1*, leading to a reduced *MKL1* expression, subsequently attenuating CAM (cell adhesion molecules) and eventually ameliorating hypoxia-induced pulmonary hypertension (Supplementary Figure 16)^30, 31^.

The high-quality genome allows us to better understand the sequences showing population-level or individual-level specificity where they are different or even absent from the human reference genome. ZF1 has more archaic-shared novel sequences than the other two Asians, consistent with a previous study proposing more archaic-shared DNAs in Tibetans than in Han Chinese^34^. The 662-bp *SCUBE2* insertion presents in both archaic and ZF1 genomes, but is absent in other Asian genomes. It is difficult to determine whether this insertion were introgressed from archaic hominids, but the association between the insertion and the lung function index (the FEV1/FVC ratio) such as the 662-bp *SCUBE2* insertion (Supplementary Table 18) suggests that these archaic-shared sequences in modern Tibetans may contribute to high altitude adaptation, in a way either selection acting on standing variants or like the reported “borrowed fitness” case of *EPAS1*^19^. Of note, *SCUBE2* plays a key role for *VEGFR2* and potentiate VEGF-induced signaling in angiogenesis. *SCUBE2* is upregulated by HIF1α at both mRNA and protein levels in lung endothelial cells^35^, providing possible mechanistic explanation of the observed association of the archaic-shared *SCUBE2* insertion with better lung functions in Tibetans.

Despite the success in charactering ZF1 SVs via analyzing TGS data, there are some limitations in this study when using the NGS data to estimate the frequency of the SVs reported from ZF1. This is mainly due to the fact that near 70% of the SVs from TGS consist of repeat elements that would cause uncertainty for short reads mapping, and most of the NGS SV detection algorithms rely on the reads mapping. Consequently, the mismapped short-reads would substantially influence the accuracy of SV detection. Given the uncertainty, we only considered those SVs with less than 70% of repeat elements and applied several stringent filtering steps to estimate the frequency difference between Tibetans and Han Chinese. This conservative strategy renders more accurate frequency estimation, while on the other hand, it might miss highly differentiated variants, especially those containing large portion of repeats. Such problem could be solved in the future when the long-read sequencing becomes cost effective for population studies.

In summary, taking advantage of long-read sequencing and next-generation mapping technologies, we *de novo* assembled a high-quality Tibetan genome and identified novel SVs, some of which might contribute to high altitude adaptation in Tibetans. Our study demonstrates the value of constructing a high-resolution reference genome of representative populations (*e.g.* native highlanders) for understanding the genetic basis of human adaptation to extreme environments as well as for future clinical applications in hypoxia-related illness.

## Methods

### ZF1 sample information

ZF1 is an adult male of native Tibetan ancestry, who has lived in Lhasa (3,680m) for more than 30 years. He is healthy (measured by physical exam and self-report), normotensive, non-anemic, normal pulmonary function and nonsmoking (by self-report). Written informed consent was obtained from ZF1. Freshly drawn blood samples were collected for DNA extraction. The protocol of this study was reviewed and approved by the Internal Review Board of Kunming Institute of Zoology, Chinese Academy of Sciences (Approval ID: SWYX-2012008) and Tibetan University (Approval ID: 2011-XZDX-001).

### Data Generation

PacBio data: high-quality genomic DNA was extracted from blood sample using the Phenol-Chloroform method and sequenced by the PacBio sequencer RSII (P6-P4 sequencing reagent).

BioNano data: according to the protocol provided by Bionano Genomics company, we obtained the high-molecular-weight DNA and constructed the high-quality sequencing library. Nt.BspQI was used for enzyme digestion. We used Irys system to analyze Bionano data.

10X Genomics data: high-molecular-weight DNA was used to construct the DNA library, and the protocol from 10X Genomic Chromium™ and Illumina Hiseq sequencer was adopted to generate long linked-reads data.

Hi-C data: adequate lymphocytes (1.5×10^7^) were extracted from fresh human peripheral blood. We constructed library by the previously published protocol and performed Illumina HiSeq PE150 sequencing to generate Hi-C data.

HiSeq XTen data: the 100× paired-end reads (150-bp) were generated using Illumina HiSeq X sequencer.

### De novo assembly

We obtained in total 24,880,404 subreads from PacBio RSII sequencer, and all these long reads were error-corrected and assembled into contigs using Falcon (v3.0) (https://github.com/PacificBiosciences/FALCON-integrate), and then polished by *Quiver*^21^. For scaffolding the contigs, we adopted two different strategies to enhance the assembling:

Strategy-1: after obtaining the assembled contigs and error-corrected long-reads, we mapped 10X Genomics linked-reads with these contigs and anchored them into preliminary scaffolds. We then generated the optical maps and the Irys BioNano platforms (Irys System, BioNano Genomics) was used in scaffolding preliminary scaffolds to elongations by the hybrid scaffold pipeline (Bionano Solve v3.1) (Supplementary Figure 2a and Supplementary Table 2).

Strategy-2: with the contigs assembled from PacBio reads, we first hybrid-assembled them into preliminary scaffolds using BioNano optical map using hybrid scaffold pipeline (same as above). Then we mapped 10X Genomics linked-reads with the preliminary scaffolds based on the supported linked-reads (Supplementary Figure 2b and Supplementary Table 3).

Next, we aligned the Hi-C reads using BWA^39^ with default parameters. Long scaffolds within each chromosomal linkage group were then assigned based on the Hi-C-based proximity-guided assembly. The original cross-linked long-distance physical interactions were then processed into paired-end sequencing libraries. First, all the reads from the Hi-C libraries were filtered by the HiC-Pro software (v2.8.1)^40^, and the paired-end reads were uniquely mapped onto the draft assembly scaffolds, which were then grouped into 24 chromosome clusters using SALSA software^22^ (Supplementary Table 22 and 23). The clustering errors were corrected by referring to GRCh38.

*PBJelly* v.15.8.24^41^ was used to close gaps of draft genomes. Briefly, all the gaps (length ≥25 bp) on the assembly were identified. Then, the long Pacbio reads were aligned to the scaffold genome using PBJelly. After read alignment, the supporting procedure was parsed by checking the multi-mapping information. After the gap-supporting sequence reads are identified, *PBJelly* assembles the reads for each gap to generate a high-quality gap-filling consensus sequence. Finally, the assembly was polished with Illumina reads by aligning the paired-end short-reads to the assembly using BWA. Picard was used to remove duplications within reads, and base-correction of the assembly was performed using *Pilon*^42^ (Supplementary Figure 2).

### Phasing the diploid assembly

We used HapCUT2^43^ and CrossStitch (https://github.com/schatzlab/crossstitch) to generate a phased genome with all the variants (SVs from PacBio, SNVs and INDELs from 10X Genomics).

### Gap closure in the human reference genome GRCh38

We closed the gaps in the human reference genome (GRCh38) by using the approach of the previous study^6^. A region consisting of continuous runs of Ns in the GRCh38 was defined as a gap. We extracted these GRCh38 gaps based on the BED file format, and the 5kb flanking sequences upstream and downstream of the gaps were mapped to the assembly by MUMmer (*nucmer -f -r -l 15 -c 25*). A gap is defined as closed only if the two flanking sequences in GRCh38 could both be aligned to the ZF1 assembly with consistent orientation, and the aligned length is over 2.5kb. The added bases were precisely counted according to the position of the two flanking sequences on the ZF1 assembly.

### Evaluation of consensus quality and sequence quality

Consensus quality of the ZF1 assembly was evaluated by comparing each chromosome with the reference genome GRCh38 using MUMmer^24^ (arguments: nucmer --mum -c 1000 -l 100; delta-filter -i 85 -l 1000 - 1).

We mapped all 100× Illumina short reads to the ZF1 assembly using the BWA-MEM^39^ module. Then we used *Picard* to mask the PCR duplicates and generated the dedup.bam file. Variants were called by the Haplotype Caller module of Genome Analysis Toolkit (GATKv3.6) (https://www.broadinstitute.org/gatk/)^44^. The SNPs and INDELs were filtered using the GATK Variant Filtration module with the following criteria, respectively: SNPs filtering: “QUAL <50; QD < 2.0; FS > 60.0; MQ < 30.0; MQRankSum < -12.5; ReadPosRankSum < -8.0; DP < 30”; INDELs filtering: “QUAL <50; QD < 2.0; FS > 200.0; ReadPosRankSum < -20.0; DP < 30”. As the previous studies described^25, 26^, we counted the total number of the homozygous SNVs (SNPs+INDELs) which represent the sites with base errors in the ZF1 assembly. The base-error rate was calculated as the number of homozygous sites divided by the total sites of the ZF1 assembly (Supplementary Table 6).

### Gene Annotation

We used GRCh38 (http://ftp.ensemblorg.ebi.ac.uk/pub/release-90/fasta/homo_sapiens/dna/) as the reference annotation panel. After performing repeat masking to ZF1 and the reference panel, we aligned ZF1 to GRCh38 by Last^45^ and generated the maf file. Then we applied CESAR2.0 (Coding Exon Structure Aware Realigner 2.0, http://github.com/hillerlab/CESAR2.0)^28^ to identify genes and the coding exons. Functional annotation for the ZF1 genes was performed using four databases: KEGG (https://www.genome.jp/kegg/), Swiss-Prot (https://www.uniprot.org/), InterPro (http://www.ebi.ac.uk/interpro/) and NR (https://www.ncbi.nlm.nih.gov/refseq/) (Supplementary Figure 5).

### Detection of SVs

For long-read PacBio data, we used mapping software NGLMR^4^ to align the error-corrected reads (‘preads’) from Falcon output to the human reference genome GRCh37. We used GRCh37 instead of GRCh38 for SV detection because the majority of the previously reported SVs were based on GRCh37, and the downstream analyses included the comparisons of these SVs among different populations. Then we used Sniffles^4^ to call SVs from the bam file and we required each variant with support from at least ten reads. For the NGS short-read Illumina data, we mapped the reads to the human reference genome GRCh37 using BWA. After sorting and removing duplicates, we used CNVnator^46^, Pindel^47^, Lumpy^48^, BreakDancer^49^ and BreakSeq2^50^ to call SVs. We further merged the results from five algorithms by MetaSV^51^. We also used PopIns^52^ to call the non-reference non-repetitive insertions from Illumina data. For BioNano data, we detected SVs by Irys *Solve (v3.1)*. For the 10X Genomics data, Long Ranger (v2.1.6) was applied to call genetic variants including SVs, SNVs and INDELs. The SNVs and INDELs were used in the phasing analysis.

### SV annotation and enrichment analysis

Annotation for ZF1 SVs were defined by VEP (http://www.ensembl.org/info/docs/tools/vep/index.html)^53^. Repeat analysis for SVs region were used by RepeatMasker v4.0.1. Function enrichment analysis was perform by DAVID v6.7^54^. In addition, we calculated the odds ratio to evaluate the enrichment of the genes affected by SVs in a set of priori candidate genes (the hypoxia regulatory genes or previously reported adaptive genes in Tibetans).

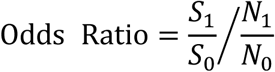

where *S1* denotes the number of SV genes presenting in the priori candidate gene list; *S0* denotes the number of SV genes absent from the priori candidate gene list; *N1* denotes the number of non-SV genes presenting in the priori candidate gene list; *N0* denotes the number of non-SV genes absent from the priori candidate gene list. The sum of *S1*, *S0*, *N1* and *N0* is the total number of genes across the genome. An odds ratio significantly above 1 (*p* < 0.05, the Chi-squared test) indicates that the SV genes are enriched in the priori gene set.

### SV genotype estimation using NGS data

We used the deletions and duplications detected by long-read sequencing platform as candidate copy number variable regions to further investigate the frequency difference in Tibetan and Han Chinese populations. The population genomic data are whole-genome sequenced (∼30X) from a previous study (38 Tibetans and 39 Han Chinese) using Illumina HiSeq X10^34^. As the short-read NGS data have bias for reads coverage at certain genomic regions^55^, we applied a stringent strategy to filter the CNV regions where we could get high quality results from NGS data. First, we filtered out the TGS CNVs where the variants could not be detected from ZF1 NGS data. Then we used CNVnator to obtain the genotype for each remaining CNV region in ZF1 sample and removed the region where the NGS genotype was inconsistent with TGS calling (consistent NGS genotype range: CNVnator genotype<1.3 for TGS deletion; 2.7<CNVnator genotype<4.3 for TGS duplication; we excluded the regions with copy number>4 due to the inaccurate estimation of high copy number variants from NGS). Next, we genotyped the remaining CNV regions for each of the 38 Tibetan and 39 Han Chinese samples using CNVnator. Based on the rounding results of the CNV genotypes, we removed the CNV regions that failed to pass the parity test^56^ in either Tibetan or Han Chinese population. V_ST_^57^ was used to measure allelic divergence between Tibetans and Han Chinese at the 1,887 CNV regions which passed all the filtering steps above.

To obtain population frequency of non-repetitive insertions, we used the assembled ZF1 genome (including the associated contigs) as reference and aligned short-reads of 38 Tibetan and 39 Han Chinese NGS data to this reference. For each insertion with sequence available reported by Sniffles, we located the positions of these sequences on the ZF1 assembly and removed the duplications using Lastz^58^ (*-- notransition --nogapped –step=20 –filter=identity:90 –filter=coverage:90*); we excluded insertions with more than 70% of repeats reported by Tandem Repeat Finder (TRF) or RepeatMasker. Next we determined the copy number (CN, e.g. 0, 1, 2..) of insertions for each sample by rounding the value of two times of the relative read-depth of the insertion. The relative read-depth was calculated as the average read depth of inserted sequence divided by the average whole genome coverage. The average read depth was calculated using SAMtools depth module. A total of 593 non-repetitive insertions were included in the analysis. Finally, to mitigate the potential batch effects from NGS data, we used a conservative way to measure the insertion frequency differentiation between Tibetan and Han Chinese by taking the minimum (mVST) of the two VST values for each insertion locus: one was directly based on CN states (VST[CN], as calculated for the CNV differentiation), and the other one (VST[norm-RD]) was based on median-normalized relative read-depth (that is, the relative read-depth divided by the median in each population; if the median equals to zero, then no normalization was performed).

We listed the top 5% of the VST and mVST for CNVs and insertions (calculated separately) in Supplementary Table 15 and 16 respectively. The 5% of the empirical VST (VST[CN]) is 0.0956 corresponding to the 98.3 percentile in the simulated null distribution (see section below).

### Simulation of SV frequency differentiation between Tibetans and Han Chinese

We employed *ms*^59^ to generate a null VST distribution to assess the SV frequency differentiation between Tibetans and Han Chinese under neutral evolution. Following previous studies^34^, we assumed that Tibetans and Han Chinese split 10,000 years ago (*T3*), and after the divergence, a bottleneck event in Tibetans occurred till 9,000 years before present (*T2*). We also considered an exponential growth of effective population size (*Ne*) for Han Chinese starting at 2,000 years before present (*T1*). We assumed the *Ne* of the Tibetan-Han Chinese common ancestry (*N1*) to be 20,000, the *Ne* at *T2* in Tibetan to be 5,000 (*N2*), and the *Ne* for present Tibetans and Han Chinese at *T3* to be 20,000 (*N3*) and 50,000 (*N4*) respectively (Supplementary Figure 18). We assumed generation time of 25 years and the mutation rate of SV to be 10^-5^ per generation^60^. The following *ms* command was used to perform the simulation: *ms 154 2000 -t 0.4 -s 1 -I 2 78 76 -g 1 458.1 -n 1 5 -n 2 2 -eg 0.002 1 0 -en 0.009 2 0.5 -ej 0.01 2 1*

### SV validation using PCR and Sanger sequencing

We genotyped the candidate SVs in ∼900 unrelated Tibetans and ∼100 unrelated Han Chinese samples using PCR and Sanger sequencing. The primers were designed by Primer Premier 5, the extended 200bp sequences were included at SV breakpoints as PCR target and sequenced using Sanger sequencing.

### Physiological traits measurement and association analysis of Tibetan populations

We collected physiological traits data and blood samples from 1,039 Tibetan volunteers who are native residents at the sampled locations, and they were from three different altitude regions in Tibet, including Lhasa (elevation: 3,658m), Bange (elevation: 4,700m) and Langkazi (elevation: 5,108m). We also sampled a Han Chinese population (n=100) from Dalian, China (elevation: 60m) as the reference. We filtered the samples based on the following criteria: 1) healthy (by physical examination and self-report); 2) normotensive, non-anemic, normal pulmonary function and non-pregnant; 3) 18≤age≤70; 4) non-smoking (by self-report). The ethnic identity was confirmed by self-claims and by report of the first language learned, and related individuals were excluded. Written informed consents were obtained from all participants.

We collected venous blood (5 ml from each individual) from subjects who fasted overnight. We measured a total of 19 physiological traits including NO: serum nitric oxide level; PAP: systolic pulmonary arterial pressure; SPO2: peripheral capillary oxygen saturation. HB: hemoglobin concentration, RBC: red blood count; HCT: hematocrit; MCV: mean red cell volume; RDW: red cell distribution width; PLT: platelets; LPC: lymphocyte count; SBP: systolic blood pressure; DBP: diastolic blood pressure; HR: heart rate; PEF: peak expiratory flow rate (L/min); MVV: Maximum Ventilatory Volume (L/min); FEF: Forced expiratory flow at 25%-75% (L/min); FEV1: Forced Expiratory Volume In 1s (%); FFR: FEV1/FVC; FVC: forced vital capacity (L). The HB concentration and other blood parameters were measured immediately using an automated hematology analyzer (Sysmex pocH-100i, Japan). SPO2 was measured at forefinger tip with a hand-held pulse oximeter (Nellcor NPB-40, CA) at rest, and the fingertip was cleaned with alcohol swab before measurement. Serum NO levels were measured by protocol as we described before^61^. For lung function test, we performed on a Microlab Spirometer 3500K, version 5.X.X Carefusion (Micro Medical Ltd., Rochester, United Kingdom) according to the ATS (American thoracic society) recommendation and the international standardized guideline^62^. Subjects were kept sitting position with a nose clip, and after two or three slow vital capacity tests, we collected the results of at least three forced vital capacity. The highest of the recorded FVC values was reported in the present study. PEF, MVV, FEF and FEV1 were measured as primary data. Stringent quality control was conducted in the entire procedure.

Genetic association analysis of physiological traits was performed using PLINK 1.07^63^. We used additive model to evaluate the association between SVs and phenotypes. Sex, age and altitude were treated as covariates for association analysis of all phenotypes. For lung functions (PEF, MVV, FEF, FEV1, FVC and FFR), we also took BMI as the covariate. To test the joint effect on association of the *MKL1*-163bp-deletion and the *SCUBE2*-662bp-insertion, we employed a joint-additive model and took the allelic status of the *MKL1* 163bp-deletion as the covariate (by plink: “*—condition*” argument). For multiple test correction, we used Benjamini & Hochberg (1995) step-up FDR control to adjust the P values using R function of *p.adjust*. We performed association test for samples from Lhasa, Bange and Langkazi separately. We performed the heterozygosity test including Cochrane’s Q statistics and I^2^ heterogeneity index (Q>0.1 and I^2^<25 for homogeneous)^64^, and found no genetic heterogeneity before pooling together the three populations.

### Non-reference sequences shared by archaic hominid and de novo assembled Asian genomes

To search for the sequences that are present in the Asian genomes (*i.e.* AK1, HX1 or ZF1) and the archaic hominids (Neanderthal or Denisovan) but absent in the human reference genome, we aligned the archaic short reads that could not be mapped to GRCh37 (reference-unmapped reads, RURs) to each of the individual genomes. We used the high-coverage Altai Neanderthal genome (∼51×) from reference^65^, and the RURs were downloaded from (http://cdna.eva.mpg.de/neandertal/altai/AltaiNeandertal/bam/unmapped_qualfail/). The high-coverage Denisovan sequencing data (∼30×) were from the literature^66^, and we processed the Denisovan raw reads following the published protocols^66^ to align to human reference GRCh37 and extracted the RURs. Then we mapped the Neanderthal and Denisovan RURs to each of the individual genomes (*i.e.* AK1, HX1 or ZF1 contigs) using *bwa aln*^39^. We treated all the archaic pair-end RURs as single-end reads during mapping. After mapping, we considered the regions with archaic RURs reaching an average depth with range between 1/3 and 1.5 folds of the genome-wide depth of the archaic reads mapped to reference human GRCh37 (*i.e.* Neanderthal: (17, 75), Denisovan: (10, 45)), as such depth range indicates that the archaic genome might likely contain one or two copies of these sequences. We further used Lastz (*--notransition - -nogapped –step=20 –filter=identity:90 –filter=coverage:90*)^58^ to align the sequences of these regions to GRCh37 and removed the sequences with high-similarity in the human reference genome. The proportion of each individual genome sharing the novel sequences with archaic hominids was calculated as the total region length with the depth falling the range above divided by the total size of the individual genome.

To obtain the positions of these sequences regarding the human reference genome, we aligned individual contigs (ZF1, HX1 and AK1) with GRCh37 using MUMmer^24^, and we only considered the sequences where their flanking positions could be determined based on GRCh37 coordinates. As shown in Supplementary Figure 17, we required both the aligned segments larger than 500-bp (c>500 & d>500) and filtered out the contigs with less than 50% coverage of alignments (a/(c+d)<0.5). We required the gap between two alignments on ZF1 contig to be larger than that on human reference genome (a>b). The region with archaic reads mapping must contain more than five reads, and the region length must be greater than half of the gap between two alignments on ZF1 contig (a’>a/2). The inserted sequences meeting all the above conditions were considered as novel sequences with clear positions on the human reference genome. As the sub-telomeric and sub-centromeric regions contain lots of repeats, we further removed the sequences located within 1Mb of telomeres or centromeres. We focused on the sequence that the archaic RUR could only align to one of the three Asian individual contigs but not the other two individual contigs. Such sequences were referred as individual-specific novel sequences shared with archaic hominids. In order to check whether these individual-specific novel sequences could be found in other modern humans other than East Asians, by using Lastz (*--notransition --nogapped –step=20 –filter=identity:95 – filter=coverage:95*), we further aligned the ZF1-specific sequences to two additional modern human *de novo* assemblies (NA12878 and NA19240 downloaded from NCBI with accession PRJNA323611)^26^, which represent European and African genomes respectively. To assess whether the ZF1-specific sequences present in non-human primates, using Lastz (*--notransition --nogapped –step=20 –filter=identity:80 – filter=coverage:80*), we aligned the sequences to the genomes of chimpanzee, gorilla and orangutan^26^.

### Estimation of EHH

To examine whether there is a selective sweep around the *SCUBE2* 622-bp insertion (Chr11: 9068607) and the *MKL1* 163-bp deletion (Chr22:40935468-40935631), we first phased the genotypes in the 1-Mb region around these variants (Chr11: 8568607-9568607 and Chr22:40435468-41435631), respectively, in the combined dataset of the whole-genome sequences of 39 Tibetan highlanders and 38 Han Chinese lowlanders, using SHAPEIT v2^67^ (r837) without any reference populations. We then calculated the EHH statistics for the focal SVs compared to the alternative alleles in these two populations, and the result was visualized using an R package *rehh*^68^ with default parameters. The ancestral allele for each locus was determined according to the ancestral sequences released by the 1000 Genomes Project.

## Data Availability

The source data can be found in the Source Data file. The PacBio sequence data, Illumina sequencing reads, the ZF1 final assembly, the phased assemblies and its annotation files are available at The Genome Sequence Archive (GSA) (http://gsa.big.ac.cn/index.jsp) under the project ID of PRJCA000936. All data can also be viewed in NODE: (http://www.biosino.org/node) by pasting the accession (OEP000207) into the text search box or through the URL: http://www.biosino.org/node/project/detail/OEP000207.

## Author contributions

B.S., X.Q., Ou., S.X. and C.C conceived and supervised the project. H.L., Y.H., L.D., Y.G., X.W. and Z.N. performed bioinformatics analysis; Ou., X.Q., C.C., Y.H., J.L., B.L., C.B., Baima., Gong., Deji., Bianba, Duoji., S.L., T.W. collected blood samples and physiological data; W.Z., Y.G., Y.H performed genotyping and sequencing validation; B.S., Y.H., H.L., L.D., X.Q. and S.X. wrote manuscript with contributions from other authors. All authors discussed the results and implications and commented on the manuscript.

## Acknowledgement

This study was supported by grants from the Strategic Priority Research Program of the Chinese Academy of Sciences (XDB13000000 to BS and SX, and XDA20040102 to XQ), the National Natural Science Foundation of China (91631306 to BS, 31671329 to XQ, 91731303, 31525014, 31771388, and 31711530221 to SX, 31460287 and 31660308 to Ou., 31601046 and 31871256 to HL), the National Key Research and Development Program of China (2017YFC1201302 to Ou. 2016YFC0906403 to SX, and 2012CB518202 to TW), the Program of Shanghai Academic Research Leader (16XD1404700 to SX), Shanghai Municipal Science and Technology Major Project (2017SHZDZX01 to SX), the UK Royal Society-Newton Advanced Fellowship (NAF\R1\191094 to SX), and the Science and Technology Commission of Shanghai Municipality (16YF1413900 to HL); the State Key Laboratory of Genetic Resources and Evolution (GREKF17-05 to JL), and the Provincial Natural Science Foundation of the Tibetan Autonomous Region (XZ2018ZR G-130 to JL) and Tibetan Fukang Hospital grant (2017-04 to JL).

## Competing interests

The authors declare no competing financial interests.

