## Supplementary Figures for "*De novo* assembly of a Tibetan genome and identification of novel structural variants associated with high altitude adaptation"

### Supplemental Data

Supplementary Figure 1 | Length distribution of raw reads and error-corrected subreads.
Supplementary Figure 2 | Overview of data generation and de novo assembly pipeline.
Supplementary Figure 3 | Comparison between the ZF1 assembly and five previous high-quality assemblies.
Supplementary Figure 4 | Dot plots of comparison between ZF1 assembly and GRCh38 assembly.
Supplementary Figure 5 | Summary of function annotation of ZF1 by different databases.
Supplementary Figure 6 | Comparison of function elements between GRCh38 and ZF1
Supplementary Figure 7 | Genome-wide distribution of structure variants of ZF1
Supplementary Figure 8 | Function enrichment of ZF1 SVs.
Supplementary Figure 9 | Manhattan plot of $V_{ST}$ between Tibetan and Han Chinese.
Supplementary Figure 10 | $mV_{ST}$ distribution of non-repetitive insertions between Tibetan and Han Chinese.
Supplementary Figure 11 | The epigenetic signals overlapped with the *MKL1* 163-bp deletion.
Supplementary Figure 12 | Genetic association analysis between three candidate SVs and multiple physiological traits in Tibetans.
Supplementary Figure 13 | Comparison of SBP and DBP among three genotypes of the *COL6A2* insertion.
Supplementary Figure 14 | Overview of insertion at *SCUBE2*.
Supplementary Figure 15 | Compared the association analysis of 622bp-insertion for *SCUBE2* and lung functions under general additive model and joint-additive model with 163bp-deletion of *MKL1*.
Supplementary Figure 16 | Schematic diagram of the putative effects of the *MKL1* 163bp-deletion on the associated phenotypes.
Supplementary Figure 17 | Illustration of novel sequence position identification on the reference genome.

Supplementary Table 1 | Summary of data generation
Supplementary Table 2 | The first version of scaffolding strategy
Supplementary Table 3 | The second version of scaffolding strategy
Supplementary Table 4 | Filled gaps at GRCh38 by ZF1
Supplementary Table 5 | Summary of ZF1 gaps
Supplementary Table 6 | Statistics of SNPs calling by NGS reads mapped to ZF1 assembly.
Supplementary Table 7 | Comparison of statistics for gene structure in ZF1
Supplementary Table 8 | Statistics for ZF1 function annotation
Supplementary Table 9 | List of the identified ZF1 SVs (GRCh37)
Supplementary Table 10 | Summary of PacBio-SVs in ZF1 supported from other platforms

Supplementary Table 11 | Repeat annotation of ZF1 SVs
Supplementary Table 12 | Functional enrichment analysis of ZF1-specific, HX1-specific and AK1-specific SVs
Supplementary Table 13 | ZF1-specific SVs located in the hypoxia-related genes
Supplementary Table 14 | ZF1-specific SVs located in the candidate genes with selective signals in Tibetans
Supplementary Table 15 | Summary of the top 5% $V_{ST}$ CNVs of ZF1
Supplementary Table 16 | Summary of the top 5% $mV_{ST}$ insertion of ZF1.
Supplementary Table 17 | Genotype frequency of the *MKL1*163bp-deletion by PCR validation and NGS calculation
Supplementary Table 18 | Results of association analysis with candidate SVs and multiple traits in Tibetans
Supplementary Table 19 | Genotype frequency of the other two SVs by PCR validation and NGS calculation
Supplementary Table 20 | ZF1-specific novel sequences shared with Neanderthals and Denisovans
Supplementary Table 21 | Genotype frequency of the *SCUBE2* 662bp-insertion by PCR validation and NGS calculation
Supplementary Table 22 | Grouping and anchoring of scaffolds with chromosomes by the Hi-C data
Supplementary Table 23 | Statistics of assembly after grouping and anchoring by the Hi-C data

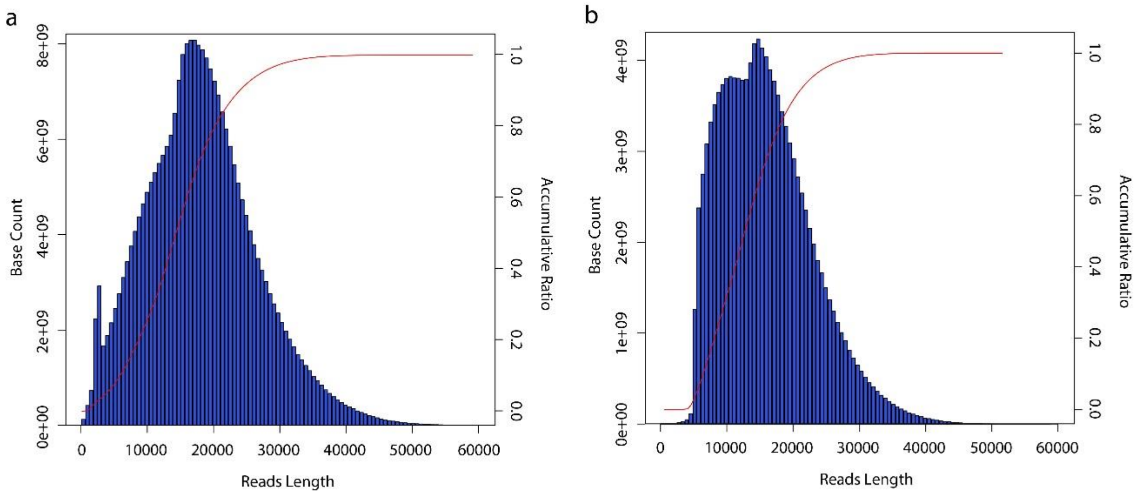


**Supplementary Figure 1 | Length distribution of raw reads and error-corrected subreads.**

a. Read length distribution of raw reads. b. Read length distribution of error-corrected subreads, which are used for de novo assembly and SV detection.

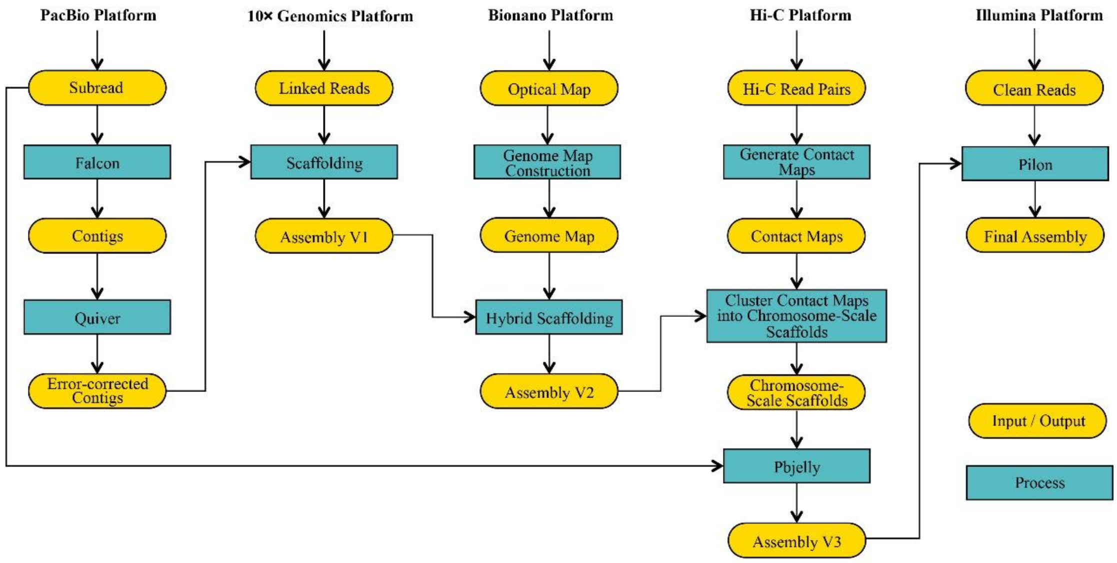


**Supplementary Figure 2 | Overview of data generation and de novo assembly pipeline.**

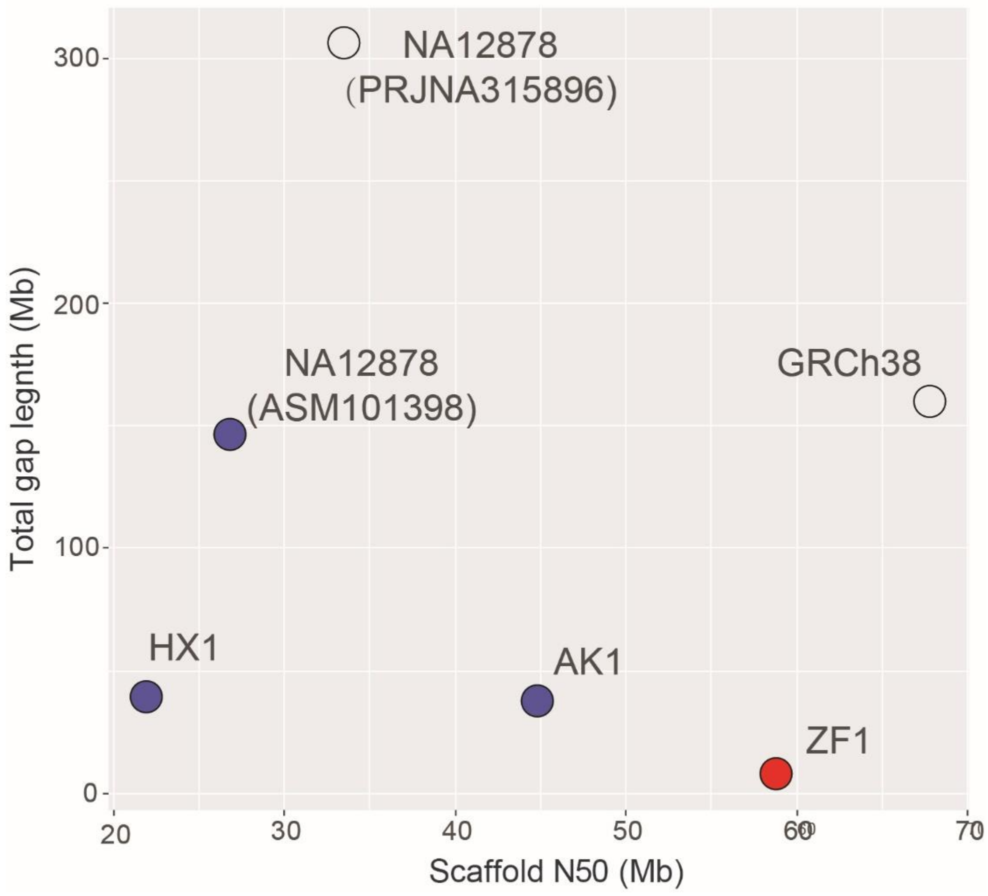


**Supplementary Figure 3 | Comparison between the ZF1 assembly and five previous high-quality assemblies.** The solid circles refer to the genome assembles by PacBio long reads, and the hollow circles refer to the genome assembles by other sequencing data without PacBio long reads.

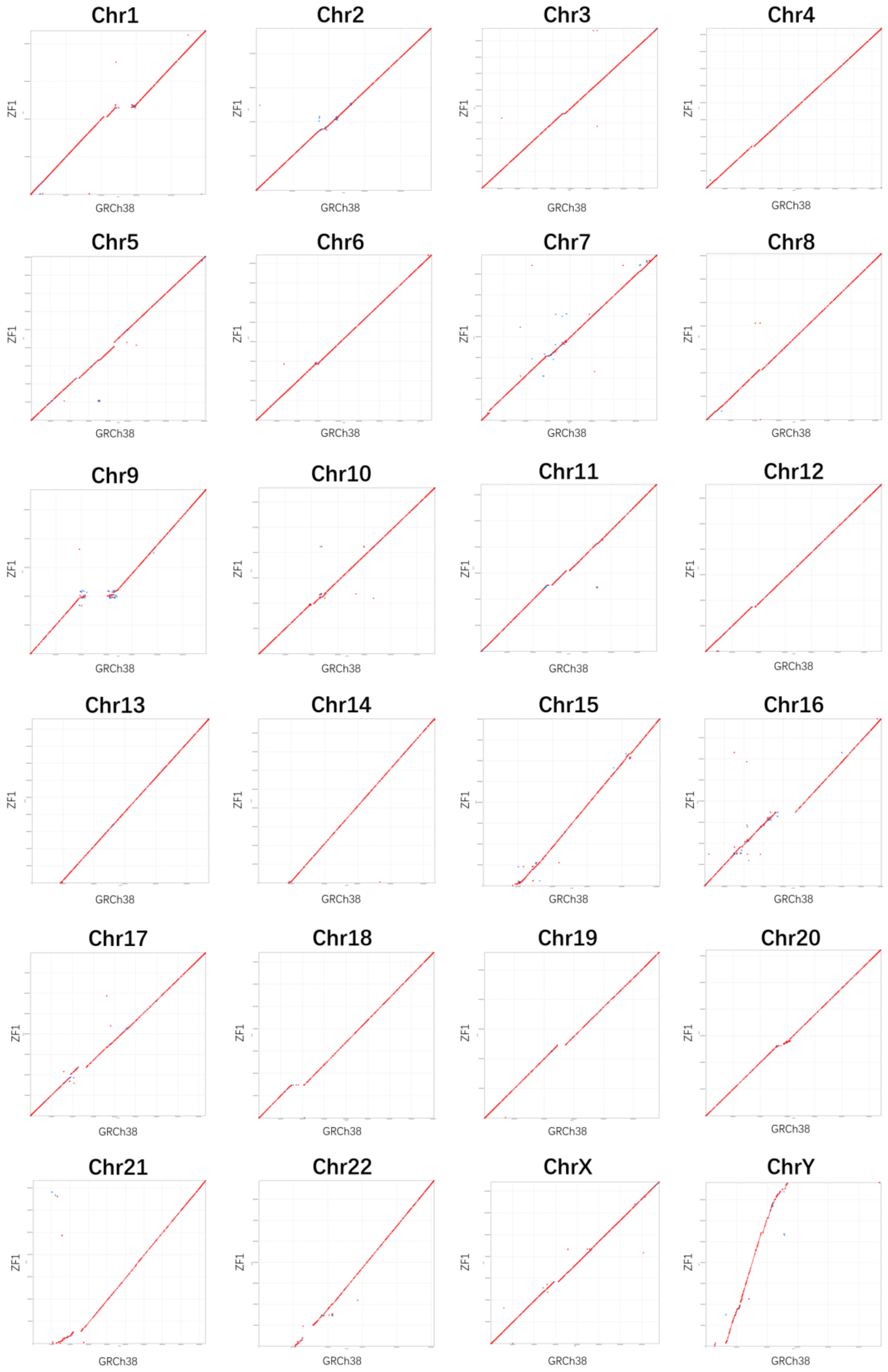


**Supplementary Figure 4 | Dot plots of comparison between ZF1 assembly and GRCh38 assembly.**

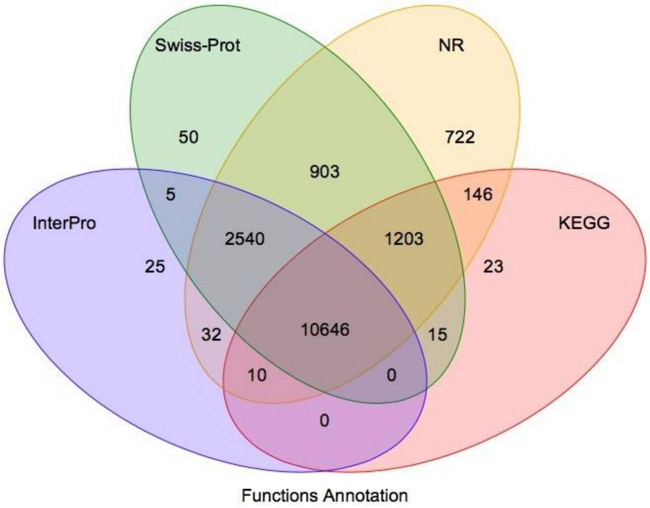


**Supplementary Figure 5 | Summary of function annotation of ZF1 by different databases.**

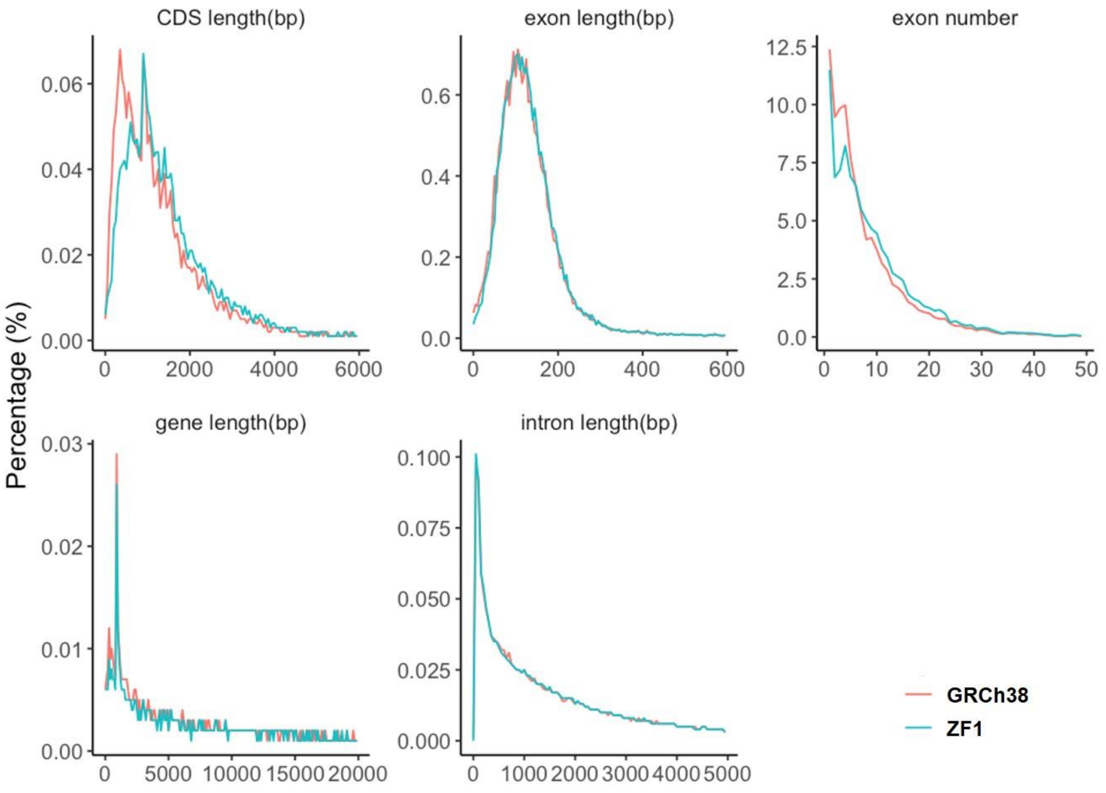


**Supplementary Figure 6 | Comparison of function elements between GRCh38 and ZF1**

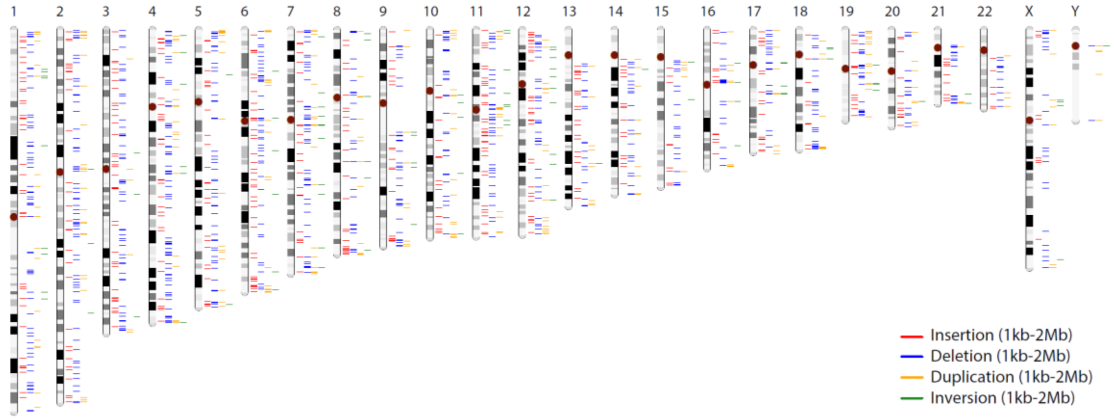


**Supplementary Figure 7 | Genome-wide distribution of large-scale structure variants of ZF1 (1Kb-2Mb)**

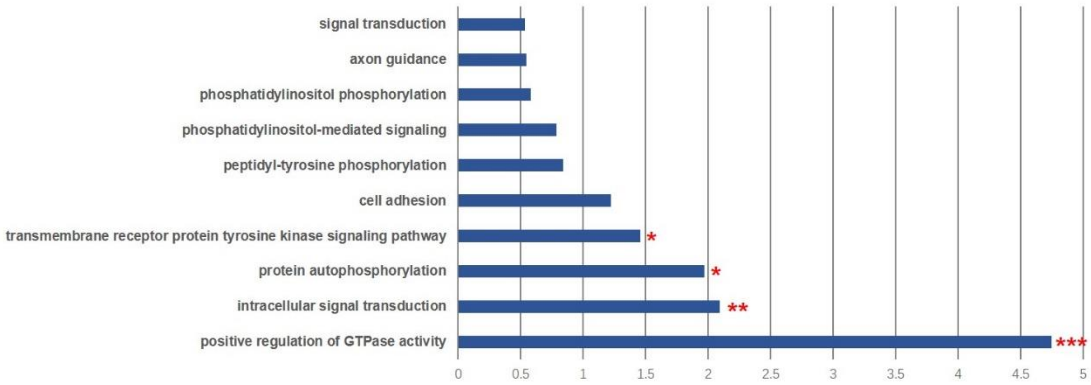


**Supplementary Figure 8 | Function enrichment of ZF1 SVs.** The significant terms were marked “*”.

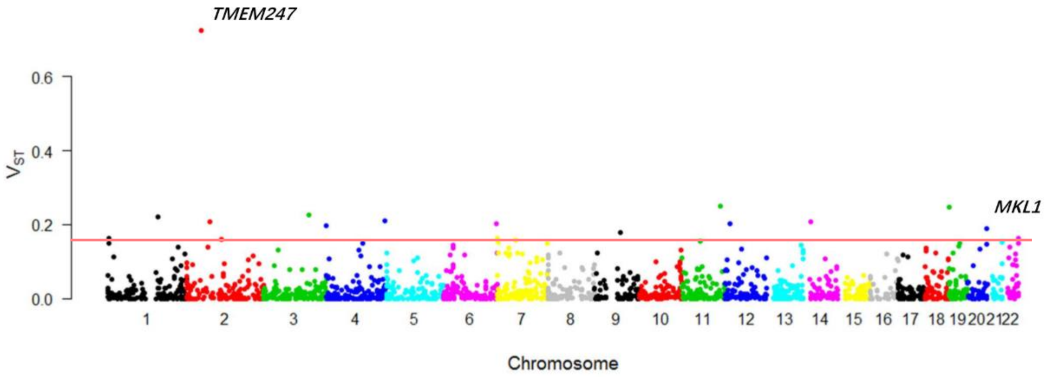


**Supplementary Figure 9 | Manhattan plot of $V_{ST}$ between Tibetan and Han Chinese.** The red line refers to the top 5% of 1887 candidate CNVs.

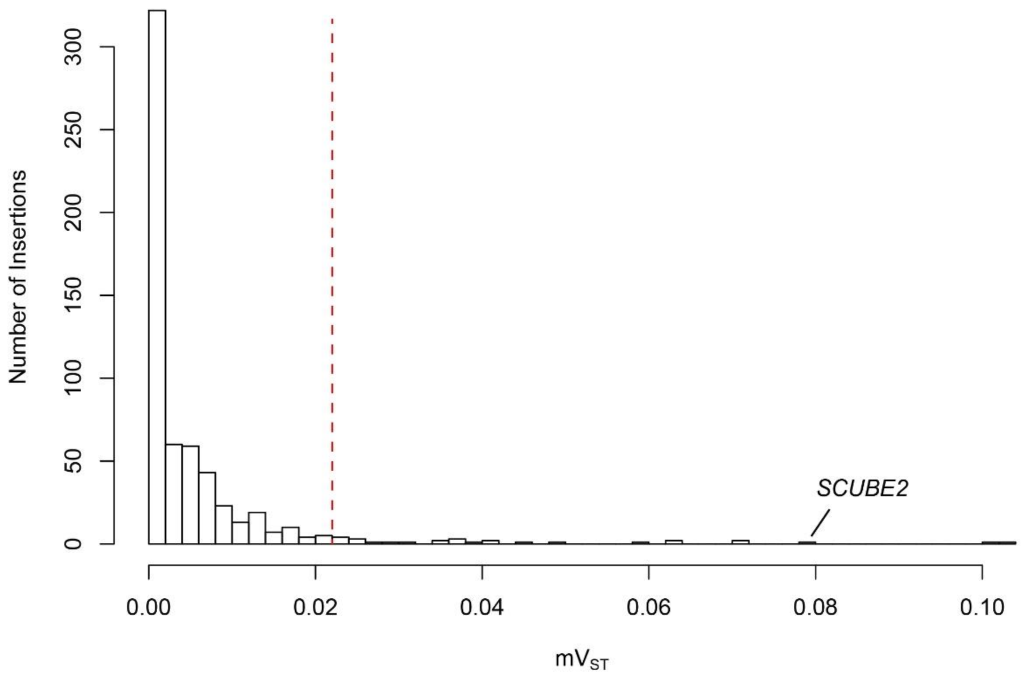


**Supplementary Figure 10 | $mV_{ST}$ distribution of non-repetitive insertions between Tibetan and Han Chinese.** The red-dashed line refers to the top 5% of 593 candidate non-repetitive insertions.

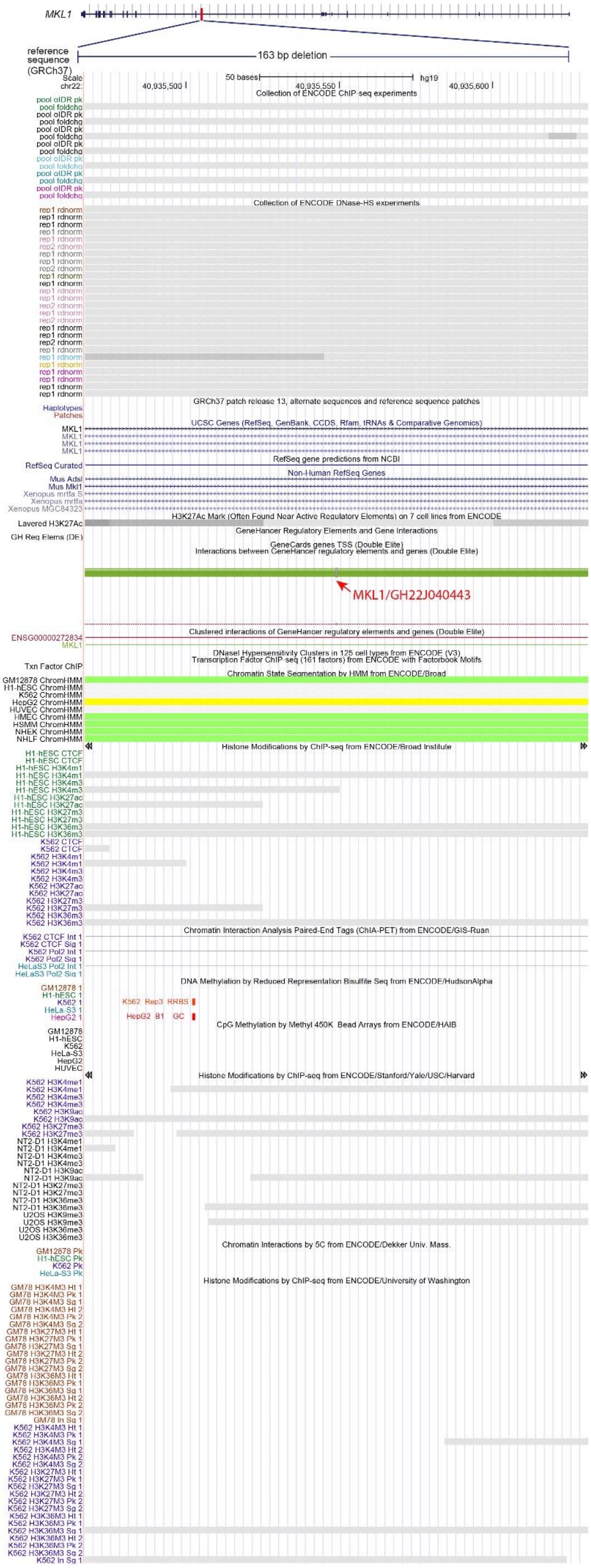


**Supplementary Figure 11 | The epigenetic signals overlapped with the *MKL1* 163-bp deletion.** The data was obtained from ENCODE at UCSC Genome Browser.

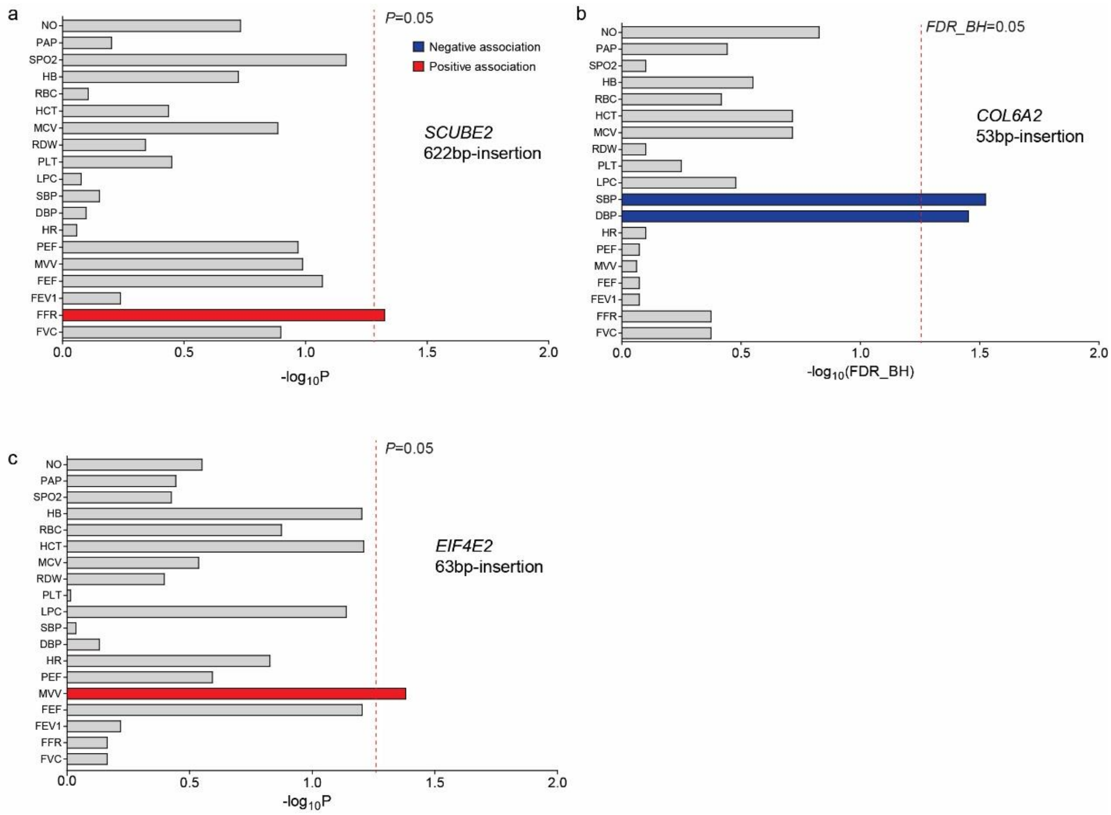


**Supplementary Figure 12 | Genetic association analysis between three candidate SVs and multiple physiological traits in Tibetans.** Dot line in red refers to significance cutoff (FDR=0.05 for b; P=0.05 for a and c), see Methods for the abbreviation of each phenotype. Blue- and red-filled histogram refers to negative and positive associated relevance, respectively.

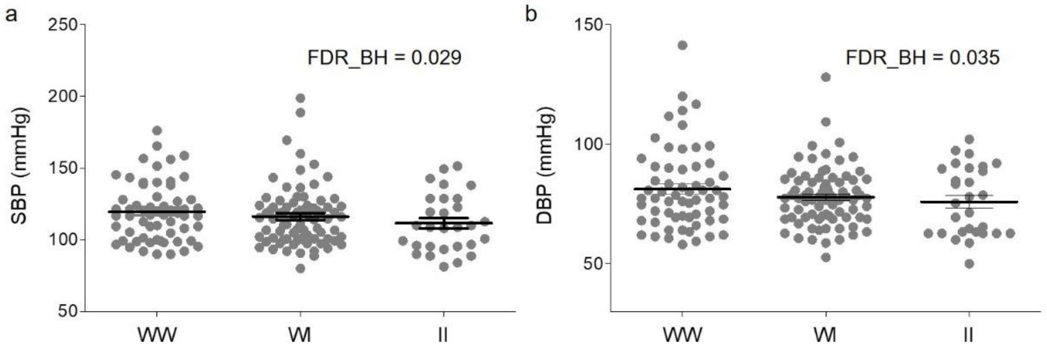


**Supplementary Figure 13 | Comparison of SBP and DBP among three genotypes of the *COL6A2* insertion.** I: 53bp insertion in *COL6A2*; W: wildtype.

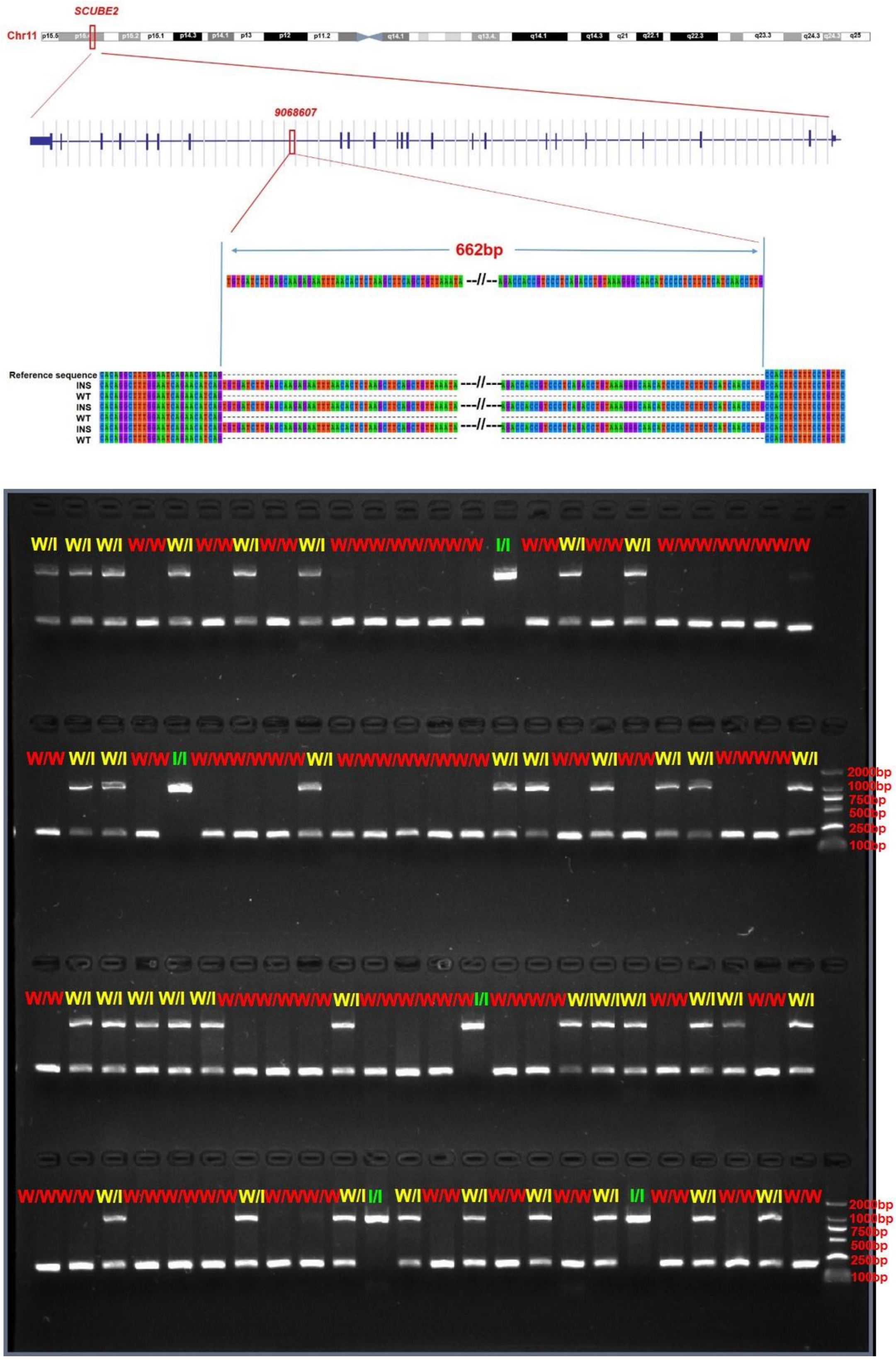


**Supplementary Figure 14 | Overview of insertion at *SCUBE2* and validation.** Sanger PCR validation of insertion region were aligned as follow (up); agarose gel electrophoresis evaluation for 662bp-insertion at SCUBE2 (down). “W”: wildtype; “I”: insertion.

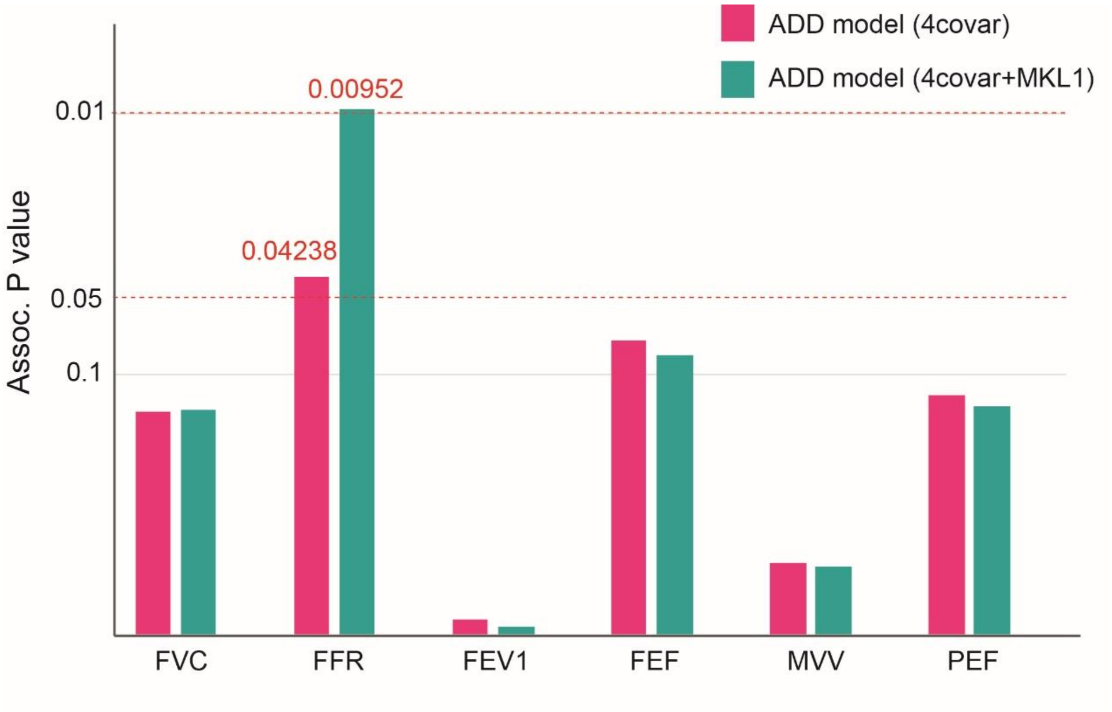


**Supplementary Figure 15 | Comparison of the P values of associations with lung functions using two different models.** The red bars indicate the P values using only the *SCUBE2* 622bp-insertion, and the green bars indicate the P values when including both the *SCUBE2* 622bp-insertion and the *MKL1* 163bp-delection.

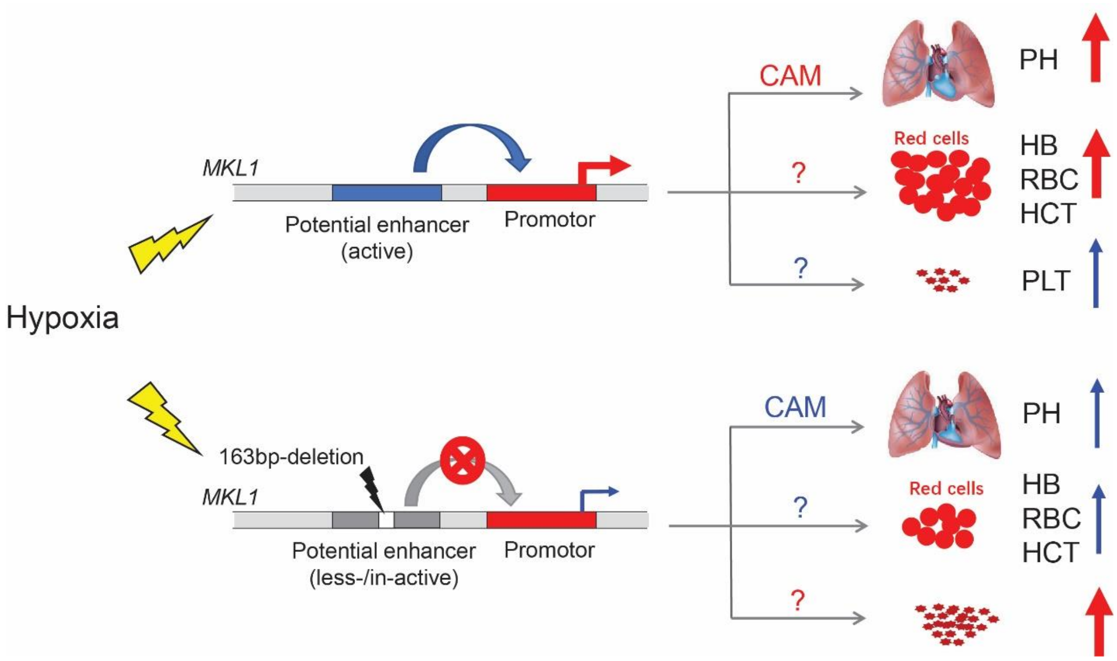


**Supplementary Figure 16 | Schematic diagram of the putative effects of the *MKL1* 163bp-deletion on the associated phenotypes.** CAM: cell adhesion molecules; PH: pulmonary hypertension; HB: hemoglobin; RBC: red blood cell; HCT: hematocrit; PLT: platelet. The regulatory pathway from CAM to PH were based on the previous studies (Chen D. et al. 2015; Yuan Z. et al. 2014).

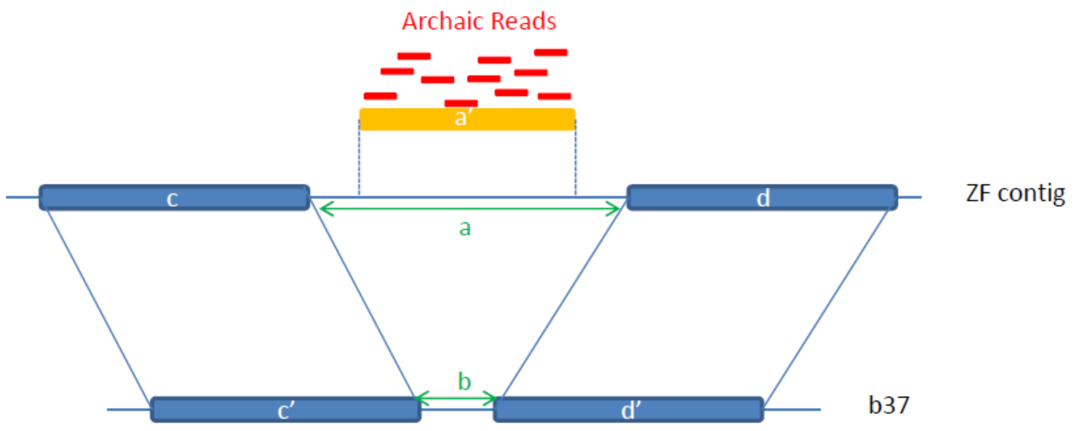


**Supplementary Figure 17 | Illustration of novel sequence position identification on the reference genome.**
The blue boxes represent the alignments of ZF1 and GRCh37 (c and c’; d and d’). The gaps between adjacent alignments are indicated in green (a, b). The red bars represent the archaic reference unmapped reads, and the orange rectangle indicates the region of these unmapped reads (a’).
